# Metabolic rewiring compensates for the loss of glutamate and aspartate biosynthesis in *Bacillus subtilis*

**DOI:** 10.1101/2023.11.10.566560

**Authors:** Mohammad Saba Yousef Mardoukhi, Johanna Rapp, Iker Irisarri, Katrin Gunka, Hannes Link, Jan Marienhagen, Jan de Vries, Jörg Stülke, Fabian M. Commichau

## Abstract

Glutamate serves as the major cellular amino group donor. In *Bacillus subtilis*, glutamate is synthesized by the combined action of the glutamine synthetase and the glutamate synthase (GOGAT). The glutamate dehydrogenases are devoted to glutamate degradation *in vivo*. To keep the cellular glutamate concentration high, the genes and the encoded enzymes involved in glutamate biosynthesis and degradation need to be tightly regulated depending on the available carbon and nitrogen sources. Serendipitously, we found that the inactivation of the *ansR* and *citG* genes encoding the repressor of the *ansAB* genes and the fumarase, respectively, enables the GOGAT-deficient *B. subtilis* mutant to synthesize glutamate via a non-canonical fumarate-based ammonium assimilation pathway. We also show that the de-repression of the *ansAB* genes is sufficient to restore aspartate prototrophy of an *aspB* aspartate transaminase mutant. Moreover, with excess nitrogen, *B. subtilis* mutants lacking fumarase activity show a growth defect that can be relieved by *aspB* overexpression, by reducing arginine uptake and by decreasing the metabolic flux through the TCA cycle. It will be interesting to investigate whether the *B. subtilis* strain using the alternative glutamate biosynthesis route can evolve in such a way that it robustly grows during nitrogen limitation and excess.

## Introduction

Glutamate is the major amino group donor in any living organism due to delivering 80 - 88% of the nitrogen for the synthesis of nitrogen-containing molecules [Wohlheuter et al., 1973; Ikeda et al., 1996; Magasanik, 1996, 2003]. Beside its role as a precursor for the synthesis of the glutamate-family amino acids like glutamine, arginine, and proline, it is also directly incorporated into proteins. To a lesser extent but still important, glutamine also functions as an amino group donor for anabolic reactions [Wohlheuter et al., 1973]. In the Gram-positive model bacterium *Bacillus subtilis*, glutamate serves as an amino group donor for at least 37 transamination reactions [Oh et al., 2007]. Therefore, it is not surprising that glutamate and glutamine are the dominating metabolites in prokaryotic and eukaryotic cells [Bennet et al., 2009; Park et al., 2016].

Glutamate also serves as a counterion for potassium ions, which are the most abundant positively charged cellular ions [Epstein, 2003]. The physiological importance of the link between glutamate and potassium was clearly demonstrated in *E. coli* that responds to an increase in medium osmolarity by accumulating potassium ions and glutamate [McLaggan et al., 1994]. Recent studies revealed that the link between glutamate and potassium homeostasis also exists in *B. subtilis* [Gundlach et al., 2018; Krüger et al., 2020, 2021]. Moreover, glutamate itself may serve as an osmoprotectant in archaea and bacteria [Csonka et al., 1994; Saum et al., 2006; Frank et al., 2021]. In *B. subtilis*, glutamate is converted to proline that serves as a compatible solute to protect the cells under hyperosmotic conditions [Zaprasis et al., 2013, 2014; Brill et al., 2011; Hoffmann et al., 2017; Bremer and Krämer, 2019; Stecker et al., 2022]. Thus, glutamate fulfils a key role in basic metabolism and the adaptation to the environmental osmolarity [Gunka and Commichau, 2012].

The enzymes catalysing the formation and degradation of glutamate link carbon to nitrogen metabolism (Figure 1A) [Commichau et al., 2006; Sonenshein, 2007]. Many organisms rely on a NADPH_2_-dependent glutamate dehydrogenase (GDH) for producing glutamate from 2-oxoglutarate and ammonium *via* reductive amination (Figure 1A) [Hudson and Daniel, 1993]. The GDH pathway was shown to be advantageous under energy limitation and at high external ammonium concentrations [Helling, 1994, 1998; Reitzer, 2003]. Alternatively, a NADPH_2_- or ferredoxin-dependent glutamate synthase (GOGAT) can be used for converting 2-oxoglutarate and glutamine into two molecules of glutamate (Figure 1A) [Suzuki and Knaff, 2005]. The glutamine required by the GOGAT is formed by the ATP-dependent glutamine synthetase (GS), a key enzyme of nitrogen metabolism that incorporates ammonium into glutamate [Kumada et al., 1993]. In contrast to the GDH-dependent ammonium assimilation, the GS-GOGAT cycle is more efficient at low ammonium concentrations because the GS has a higher affinity for ammonium than the GDH [Helling, 1994, 1998; Reitzer, 2003].

**Fig. 1.**
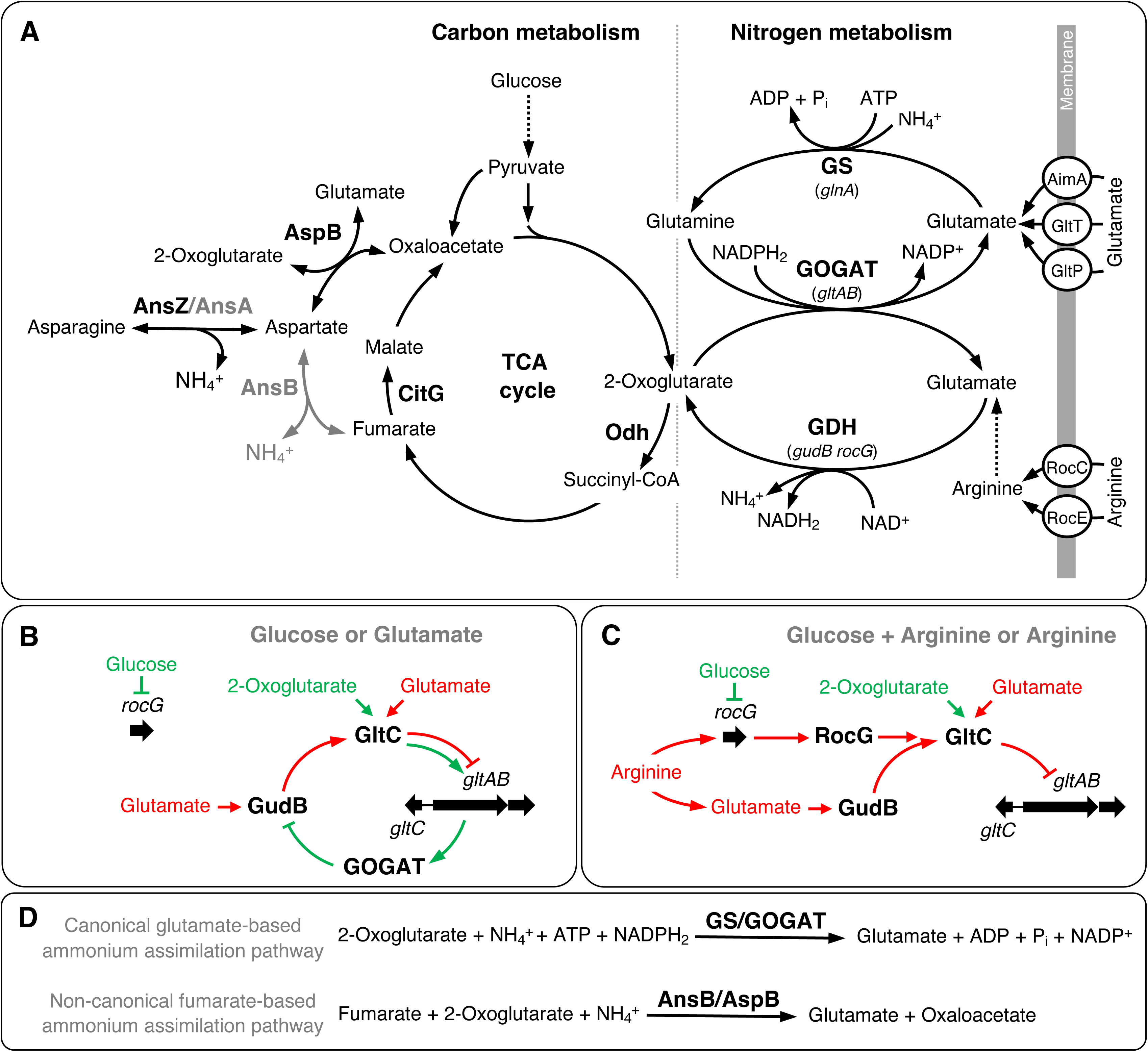
Links between carbon and nitrogen metabolism and regulation of glutamate biosynthesis in *B. subtilis.* **A.** Aspartate and glutamate metabolism and transporters for arginine and glutamate. AimA, GltP and GltT, glutamate transporters; RocC and RocE, arginine permeases; AnsA and AnsZ, asparaginases; AnsB, aspartase; AspB, aspartate transaminase; CitG, fumarase; Odh. 2-oxoglutarate dehydrogenase enzyme complex; GDH, glutamate dehydrogenase encoded by *gudB* or *rocG*; GOGAT, glutamate synthase encoded by *gltAB*; GS, glutamine synthetase encoded by *glnA*; TCA, tricarboxylic acid. **B.** Glucose- and glutamate-dependent induction and repression, respectively, of the *gltAB* glutamate synthase genes. **C.** Arginine-dependent repression of the *gltAB* glutamate synthase genes. **D.** Glutamate- and fumarate-based ammonium assimilation pathways.

We are interested in glutamate metabolism of *B. subtilis* that possesses the GS-GOGAT cycle and two GDHs [Bohannon and Sonenshein, 1989; Belitsky and Sonenshein, 1998]. Unlike *E. coli*, the NAD^+^-dependent GDHs RocG and GudB of *B. subtilis* are strictly devoted to glutamate degradation, most probably because of their low affinity for ammonium [Commichau et al., 2008; Gunka et al., 2010]. While the *gudB* gene is constitutively expressed, the *rocG* gene is subject to complex regulation by signals derived from carbon and nitrogen metabolism [Gunka and Commichau, 2012]. In the presence of carbon sources that exert carbon catabolite repression, the pleiotropic transcription factor CcpA prevents the transcription of the *rocG* gene [Belitsky et al., 2004; Choi and Saier, 2005]. Under these conditions, the LysR-type family transcription factor GltC activates the expression of the *gltAB* GOGAT genes (Figure 1B) [Bohannon and Sonenshein, 1989; Wacker et al., 2003; Belitsky and Sonenshein, 2004; Commichau et al., 2006a]. The binding of GltC to the *P_gltAB_* promoter is stimulated by 2-oxoglutarate (Figure 1B) [Picossi et al., 2007]. Recently, it has been shown that the synthesized GOGAT GltAB binds to and inactivates the GDH GudB to prevent the degradation of glutamate that is required by the bacteria to achieve high growth rates (Figure 1B) [Jayaraman et al., 2022; Hartmann, 2022]. The GOGAT can now be assigned to the group of moonlighting proteins because it fulfils two functions: glutamate synthesis and inactivation of the GDH GudB [Liu and Jeffery, 2020]. Undomesticated strains of *B. subtilis* like NCIB 3610 can also use glutamate as the single source of carbon and nitrogen. Under these conditions, the catalytically active GDH GudB binds to and prevents GltC from activating the transcription of the *gltAB* genes (Figure 1B) [Stannek et al., 2015; Noda-Garcia et al., 2017]. A recent study suggests that the GDH GudB may convert the transcription activator GltC to a repressor of the *gltAB* genes, depending on the availability of glutamate [Dormeyer et al., 2019]. This regulatory system allows *B. subtilis* to grow under fluctuating glutamate concentrations [Jayaraman et al., 2022]. In the presence of arginine, which is converted via ornithine to glutamate, the DNA binding transcription factors AhrC and RocR, of which the latter is triggered by ornithine, activate the transcription of the *rocG* gene and the encoded GDH RocG [Calogero et al., 1994; Gardan et al., 1995; 1997; Miller et al., 1997; Warneke et al., 2023]. Like the GDH GudB, RocG can also inactivate the GltC protein [Commichau et al., 2007a, 2007b; Stannek et al., 2015]. The arginine-dependent induction of the *rocG* gene dominates the glucose-dependent negative regulation exerted by CcpA [Commichau et al., 2007a, 2007b; Stannek et al., 2015]. Thus, the GDH RocG only comes into play when amino acids of the glutamate family (e.g., arginine) are available in the environment in high amounts [Stannek et al., 2015]. The redundancy of the GDH-encoding genes and their differential regulation probably provide the bacteria with a selective growth advantage when high-level GDH activity is required. The domesticated *B. subtilis* strains 160, 166 and 168 harbour the cryptic *gudB^CR^* allele encoding an inactive GDH due to a tandem repeat in the open reading frame [Belitsky and Sonenshein, 1998; Zeigler et al., 2003; Gunka et al., 2012]. It has been shown that the spontaneous decryptification of the *gudB^CR^* allele allow the bacteria to utilize amino acids of the glutamate family more efficient [Belitsky and Sonenshein, 1998; Gunka et al., 2012, 2013]. Thus, the genetic makeup of the laboratory strain 168 does not reflect the natural situation in undomesticated *B. subtilis* strains. In fact, GudB and RocG must be considered as the major and minor GDHs, respectively, of *B. subtilis*, [Stannek et al., 2015]. However, both GDHs are trigger enzymes, which are a subclass of moonlighting proteins that are active in metabolism and in controlling gene expression [Commichau and Stülke, 2008]. *B. subtilis* can also take up glutamate from the environment *via* the glutamate transporters AimA and GltT [Zaprasis et al., 2015; Krüger et al., 2021]. Moreover, GltT and GltP mediate the transport of the herbicides glyphosate and glufosinate into the cell [Wicke et al., 2019; Schwedt et al., 2023].

Here, we show that the glutamate and aspartate auxotrophies of *B. subtilis* mutants lacking the GOGAT and aspartate transaminase AspB, respectively, are relieved by *ansR* and *citG* mutations that open an alternative entry point for ammonium via the reaction that is catalyzed by the L-aspartase AnsB. Genetic and cultivation experiments as well as metabolome analyses confirmed that metabolic rewiring of central metabolism compensates for the loss of glutamate and aspartate biosynthesis in *gltAB* and *aspB* mutants, respectively. We also observed that the *B. subtilis citG* mutants lacking fumarase activity showed a growth defect when excess nitrogen was available. This finding underlines the importance of the tight regulation of glutamate biosynthesis and degradation, which is the case for the canonical glutamate-based ammonium assimilation pathway in *B. subtilis*.

## Results

### Mutations in the *ansR* and *citG* genes relieve glutamate auxotrophy of a *gltAB* mutant

Serendipitously, we observed that the *B. subtilis gltAB* mutant BP261 lacking the GOGAT formed single colonies on CGXII minimal medium plates during incubation for 5 days at room temperature (Figure 2A). CGXII medium that is commonly used for growth and maintenance of *Corynebacterium glutamicum*, contains glucose and ammonium/urea as sources of carbon and nitrogen, respectively [Keilhauer et al., 1993]. Single colonies of two potential *gltAB* suppressor mutants designated as BP364 (M1) and BP365 (M2) as well as the parental strain BP261 (*gltAB*) as a control were propagated on CGXII plates supplemented with tetracycline (*tet*). In contrast to the parental strain, the suppressor mutants grew after 48 h of incubation at 37°C (Figure S1A). Next, we verified the replacement of the *gltAB* gene by the *tet* resistance gene in two suppressors by PCR (Figure S1B). To assess whether the suppressor mutations are genetically linked to the *gltAB* locus, we isolated the chromosomal DNAs of the two mutants and transformed the wild-type strain SP1. Subsequent growth experiments revealed that the transformants were unable to grow on CGXII plates lacking glutamate. Thus, the mutation(s) in the mutants M1 and M2 are not genetically linked to the *gltAB* locus.

**Fig. 2.**
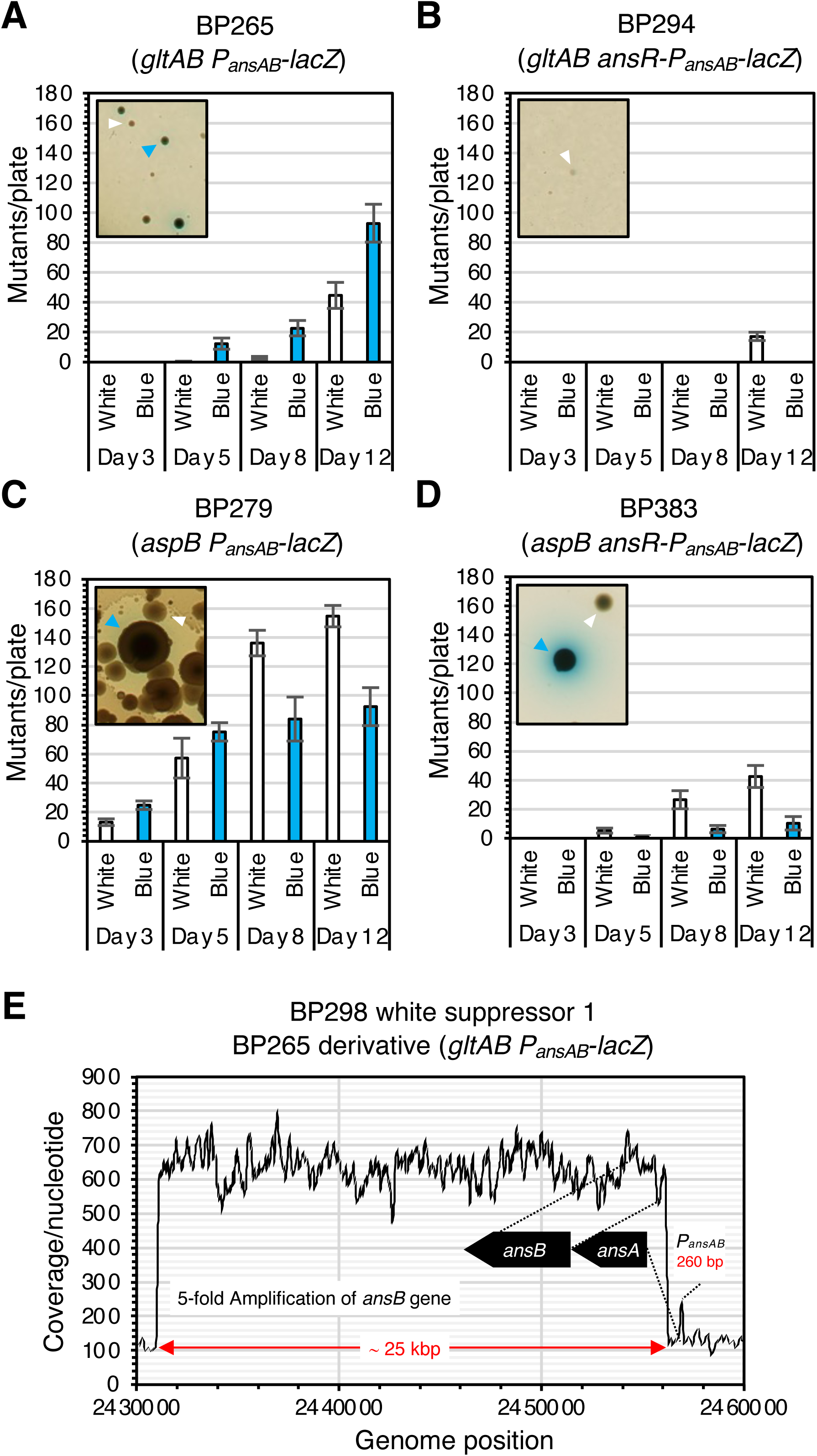
Genomic adaptation of *B. subtilis aspB* and *gltAB* mutants abrogates aspartate and glutamate auxotrophy. **A.** Formation of blue and white suppressor mutants (indicated by blue and white arrows) by the strains BP265 (*gltAB P_ansAB_-lacZ*), BP294 (*gltAB ansR-P_ansAB_-lacZ*), BP279 (*aspB P_ansAB_-lacZ*) and BP383 (*aspB ansR-P_ansAB_-lacZ*) harbouring a translational *ansAB* promoter-*lacZ* fusion during growth under selection on CGXII plates. Strains BP294 and BP383 carry an additional *ansR* copy in the *amyE* locus. The activity of the *P_ansAB_* promoter was visualized by adding the chromogenic substrate X-Gal to the plates. **B.** Read coverage along the chromosomal segment ranging from 2.430.000 to 24.600.000 bp. Based on the average coverage of the amplified region and of the remaining genome it can be inferred that five copies of the 25 kbp long region containing the entire *ansB* gene are present in the suppressor BP298.

Next, we identified mutations in the mutants M1 and M2 by whole-genome sequencing. As shown in Table 1, the sequencing analyses revealed that both mutants had acquired mutations affecting the *ansR* and *citG* genes. The *ansR* gene encodes the transcriptional repressor AnsR of the *ansAB* L-asparaginase and L-aspartase genes that are required for asparagine and aspartate degradation [Sun and Setlow, 1991, 1993; Fisher and Wray, 2002]. The single nucleotide exchanges in the *ansR* alleles of M1 and M2 would cause the amino acid replacements C57R and L101P, respectively, in AnsR. The AnsR variants are likely impaired in their ability to bind to the *P_ansAB_* promoter because the residues C57 and L101 are close to the helix-turn-helix motif and in the dimerization domain, respectively (Figure S1C). The *citG* gene codes for the fumarase CitG that catalyzes the conversion of fumarate to malate in the tricarboxylic acid (TCA) cycle (Figure 1A) [Feavers et al., 1988; Moir et al., 1984; Miles and Guest, 1985]. While the mutant M1 does not synthesize the fumarase CitG because a 10.7 kbp-long region (coordinates 3381391 to 3392113) including a big part of the *citG* gene was deleted, a single nucleotide insertion in the *citG* gene in mutant M2 would truncate the encoded protein (Table 1). Thus, two genomic alterations are sufficient to relieve glutamate auxotrophy of the *B. subtilis gltAB* mutant. Obviously, the reduced DNA-binding activity of AnsR causes de-repression of the *ansAB* genes, thereby allowing the GOGAT-deficient bacteria to synthesize glutamate from ammonium and fumarate via the L-aspartase AnsB and aspartate transaminase AspB that catalyze reversible reactions (Figures 1A and 1D) [Viola, 2000]. Moreover, the lack of CitG fumarase activity likely enhances the metabolic flux from the TCA cycle to the GOGAT bypass.

**Table 1:** Identified mutations in the *gltAB* and *aspB* suppressor mutants.

| **Table 1: Identified mutations in the *gltAB* and *aspB* suppressor mutants** |  |  |  |  |  |
| --- | --- | --- | --- | --- | --- |
| **Mutations relieving glutamate auxotrophy of the strains BP261 (*gltAB*) and BP265 (*gltAB P<sub>ansAB-lacZ</sub>*)** |  |  |  |  |  |
| **Strain** | **Mutant** | **Parental strain** | **Phenotype** | **Affected gene, mutations** | **Amino acid exchanges, effect on the protein** |
| BP364 | M1<sup>A</sup> | BP261 (*gltAB*) | Growth on CGXII | *ansR* T302C, 10.7 kbp deletion including *citG* | AnsR L101P, CitG not synthesized |
| BP365 | M2<sup>A</sup> | BP261 (*gltAB*) | Growth on CGXII | *ansR* T169C, *citG* Δ62G | AnsR C57R, CitG ΔC43 |
| BP281 | S1B<sup>B</sup> | BP265 (*gltAB P<sub>ansAB-lacZ</sub>*) | Growth on CGXII, blue colonies | *ansR* +A36, *citG* +G917 | AnsR ΔC11, CitG ΔC310 |
| BP282 | S2B<sup>B</sup> | BP265 (*gltAB P<sub>ansAB-lacZ</sub>*) | Growth on CGXII, blue colonies | *ansR* Δ2456934-2457298 | AnsR ΔC16 |
| BP283 | S3B<sup>B</sup> | BP265 (*gltAB P<sub>ansAB-lacZ</sub>*) | Growth on CGXII, blue colonies | *ansR* +A36, *citG* C1226A | AnsR ΔC11, CitG A409E |
| BP284 | S4B<sup>B</sup> | BP265 (*gltAB P<sub>ansAB-lacZ</sub>*) | Growth on CGXII, blue colonies | *ansR* +A36 | AnsR ΔC11 |
| BP285 | S5B<sup>B</sup> | BP265 (*gltAB P<sub>ansAB-lacZ</sub>*) | Growth on CGXII, blue colonies | *ansR* +A36 | AnsR ΔC11 |
| BP286 | S6B<sup>B</sup> | BP265 (*gltAB P<sub>ansAB-lacZ</sub>*) | Growth on CGXII, blue colonies | *ansR* +A36 | AnsR ΔC11 |
| BP287 | S7B<sup>B</sup> | BP265 (*gltAB P<sub>ansAB-lacZ</sub>*) | Growth on CGXII, blue colonies | *P<sub>ansAB</sub>* (G-10A), *citG* T876A | Enhanced *ansAB* expression, CitG D292E |
| BP288 | S8B<sup>B</sup> | BP265 (*gltAB P<sub>ansAB-lacZ</sub>*) | Growth on CGXII, blue colonies | *ansR* +A36 | AnsR ΔC11 |
| BP289 | S9B<sup>B</sup> | BP265 (*gltAB P<sub>ansAB-lacZ</sub>*) | Growth on CGXII, blue colonies | *ansR* +A36 *citG*, Δ3390211-3390276 | AnsR ΔC11, CitG not synthesized |
| BP290 | S10B<sup>B</sup> | BP265 (*gltAB P<sub>ansAB-lacZ</sub>*) | Growth on CGXII, blue colonies | *ansR* +A36 | AnsR ΔC11 |
| BP298 | S1W<sup>A</sup> | BP265 (*gltAB P<sub>ansAB-lacZ</sub>*) | Growth on CGXII, white colonies | *citG* A1033T, 25.2 kbp amplification including *ansB* | CitG I345F, increased AnsB synthesis |
| **Mutations relieving aspartate and asparagine auxotrophy of the strain BP279 (*aspB P<sub>ansAB-lacZ</sub>*)** |  |  |  |  |  |
| BP366 | S1B<sup>B</sup> | BP279 (*aspB P<sub>ansAB-lacZ</sub>*) | Growth on CGXII, blue colonies | *ansR* C171A | AnsR ΔC56 |
| BP367 | S2B<sup>B</sup> | BP279 (*aspB P<sub>ansAB-lacZ</sub>*) | Growth on CGXII, blue colonies | *ansR* G3A | AnsR M1I |
| BP368 | S3B<sup>B</sup> | BP279 (*aspB P<sub>ansAB-lacZ</sub>*) | Growth on CGXII, blue colonies | *ansR* G94C | AnsR A32P |
| BP296 | S1W<sup>A</sup> | BP279 (*aspB*<br>*P<sub>ansAB</sub>-lacZ*) | Growth on CGXII,<br>white colonies | *ansA* T43A, *citG* C686T | AnsA S15T, CitG A229V |
| BP297 | S2W<sup>A</sup> | BP279 (*aspB*<br>*P<sub>ansAB</sub>-lacZ*) | Growth on CGXII,<br>white colonies | *ansA* T43A, *citG* C312G | AnsA S15T, CitG N104K |

<sup>A</sup>Mutations were identified by genome sequencing.
<sup>B</sup>Mutations were identified by Sanger sequencing with primer pairs SM2/SM18 and SM3/SM4 for *P<sub>ansAB</sub>-ansR* and *citG*, respectively.

### A selection and screening system to distinguish *gltAB* suppressor mutants

To assess whether the inactivation of *ansR* is a prerequisite for restoring glutamate prototrophy of the *gltAB* mutant, we established a selection and screening system. For this purpose, we introduced a translational *P_ansAB_-lacZ* into the *gltAB* mutant and propagated the strain BP265 (*gltAB P_ansAB_-lacZ*) on CGXII agar plates supplemented with the chromogenic substrate X-Gal. Blue suppressor mutants would indicate the accumulation of loss-of-function mutations in the *ansR* gene. By contrast, white mutants would either indicate the inactivation of the *citG* gene or the selective amplification of a genomic locus containing the *ansAB* operon as previously observed in other suppressor analyses [Dormeyer et al., 2017; Wicke et al., 2019; Richts et al., 2021]. As shown in Figure 2A, several suppressor mutants appeared on the CGXII-X-Gal plate after incubation for 3 days at 37°C. Sanger sequencing of the *ansR* and *citG* genes in ten isolated mutants revealed that all mutants had acquired mutations in the *ansR* or in ribosome-binding site of the *P_ansR_* promoter (Figure S1D), and in four of the mutants the *citG* gene was also affected (Table 1) (Figure S1E). Thus, the de-repression of the *ansAB* genes either by mutational inactivation of *ansR* or by mutations interfering with *ansR* expression is important for allowing the *gltAB* suppressors to synthesize glutamate via the AnsB/AspB-dependent route (Figure 1A). Moreover, mutations that inactivate *citG* or lowering the activity of the fumarase probably contribute to redirect the metabolic flux to the aspartate branch.

Further incubation of the CGXII-X-Gal plates carrying the blue *gltAB* suppressor mutants for up to 8 days resulted in the emergence of additional white suppressors (Figure 2A). Illumina sequencing of one arbitrarily chosen white mutant BP298 revealed that the strain had amplified a 25.3 kbp-long genomic segment causing a 5-fold increased dosage of the *ansAB* operon and its surrounding genes (Figure 2E. Thus, the glutamate auxotrophy of the strain BP265 (*gltAB P_ansAB_-lacZ*) can also be relieved by enhancing the dosage of the *ansAB* genes that are not expressed in the parental strain. We also observed that the *gltAB* mutant BP294 (*gltAB ansR-P_ansAB_-lacZ*) containing the native *ansR* gene and a second *ansR* copy in the *amyE* locus formed much less and only white mutants (Figure 2B). Thus, *ansR* loss-of-function mutations are indeed crucial for relieving glutamate auxotrophy of a *gltAB* mutant. Finally, we tested whether the strain BP280 (*gltAB ansAB*) lacking GOGAT and AnsAB was able to form suppressors on CGXII plates (Figure 2C). Like the *gltAB* mutant, also the *aspB* BP294 (*aspB ansR-P_ansAB_-lacZ*) containing two *ansR* copies formed less and mainly white mutants (Figure 2D). Moreover, we propagated the strain BP280 (*gltAB ansAB*) on CGXII plates. However, this strain did not form suppressor mutants (data not shown). This finding suggests that there is no other way to bypass the reactions catalysed by GOGAT and AnsB/AspB.

### Inactivation of *ansR* relieves aspartate auxotrophy of an *aspB* mutant

*B. subtilis* synthesizes aspartate via the L-aspartate transaminase AspB that converts oxaloacetate and glutamate to 2-oxoglutarate and aspartate (Figure 1A). Previously, it has been shown that an *aspB* mutant is auxotrophic for aspartate and asparagine [Zhao et al., 2018]. The study by Zhao et al. also revealed a slightly heterogenous colony morphology of a L-aspartase transaminase-deficient *B. subtilis* strain, indicating genetic instability of the mutant during growth on aspartate-limited plates (Figure 2C in Zhao et al., 2018). Therefore, we hypothesized that the aspartate prototrophy of the *aspB* mutant could be restored by the inactivation of *ansR*, facilitating aspartate synthesis from ammonium and fumarate via the reversible L-aspartase AnsB (Figure 1A) [Viola, 2000]. To test this idea, we propagated the *aspB* mutant BP279 (*aspB P_ansAB_-lacZ*) on CGXII-X-Gal plates. Again, the translational *P_ansAB_-lacZ* fusion was used to facilitate the discrimination between different mutant classes. As observed with the *gltAB* mutant, several blue colonies appeared on the plates after incubation for 3 days at 37°C (Figure 2A). Sanger sequencing of *ansR* in the three isolated *aspB* suppressors BP366, BP367 and BP368 indeed revealed that the mutants had acquired loss-of-function mutations in the *ansR* gene that would inactivate the encoded repressor (Table 1) (Figure S1C). Moreover, as observed with the *gltAB* mutant, the *aspB* mutant BP383 (*aspB ansR-P_ansAB_-lacZ*) containing two *ansR* copies formed much less and only white mutations (Figure 2A). In contrast to the *gltAB* mutant strain BP265, the *aspB* mutant BP279 produced more suppressors, including significantly more that showed a white phenotype (Figure 2A). The genome sequencing analysis of the two isolated white *aspB* suppressors BP296 and BP297 revealed that both mutants had acquired mutations in the *citG* gene that probably reduce the activity of CitG (Figure S1E) (Table 1). Both mutants also carry a mutation in the *ansA* gene (Table 1). It is tempting to speculate that the S15T replacement in AnsA reduces asparagine production from aspartate, which is needed by the *aspB* mutant (Figure 1A). To conclude, the aspartate auxotrophy of the *aspB* mutant can be alleviated by mutations affecting the activity of AnsR and CitG.

### Characterization of mutants synthesizing glutamate and aspartate via AnsB

To assess the growth behaviour of *gltAB* mutants synthesizing glutamate and aspartate via the L-aspartase AnsB, we deleted the *ansR* and *citG* genes individually and in combination in the strain BP265 (*gltAB*). In the strain BP279 (*aspB*), we only tested the effect of *ansR* deletion because it seems to be sufficient to completely relieve aspartate auxotrophy (Table 1). Next, we evaluated the growth of the strains BP273 (*gltAB ansR*), BP274 (*gltAB citG*), BP276 (*gltAB ansR citG*), and BP292 (*aspB ansR*) on CGXII agar in the absence and in the presence of either glutamate, aspartate, or asparagine. The parental strains BP265 (*gltAB*) and BP279 (*aspB*) served as controls. In liquid medium, the growth of the parental and the reconstituted strains was assessed. As shown in Figure 3A and 3B, except for the strain BP279 (*aspB*), which cannot convert glutamate to aspartate, the remaining strains grew on plates and in liquid medium supplemented with glutamate as an additional nitrogen source. Moreover, as expected, aspartate supported growth of all strains. By contrast, on agar plates supplemented with asparagine, the strains BP274 and BP276 lacking the *citG* fumarase gene showed a growth defect, which was not the case for strain BP276 in liquid medium (Figure 3A and 3B). Moreover, only the reconstituted strains BP276 (*gltAB ansR citG*) and BP292 (*aspB ansR*) showed significant growth on plates and in liquid medium containing ammonium as the single source of nitrogen. Thus, the auxotrophy for glutamate and aspartate of the *gltAB* and *aspB* mutants can be relieved by the inactivation of the *ansR citG* and *ansR* genes, respectively.

**Fig. 3.**
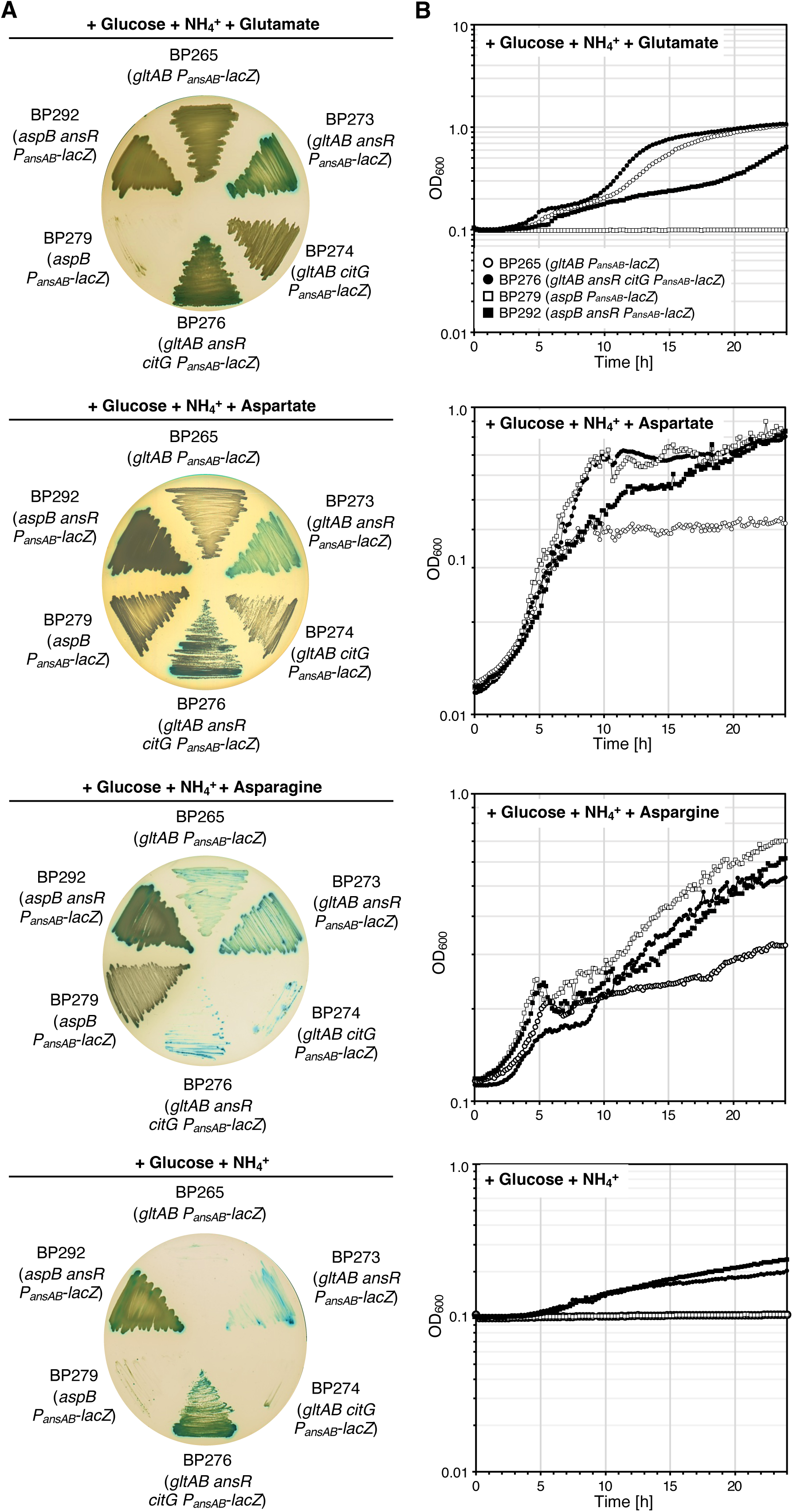
Characterization of *B. subtilis* strains synthesizing aspartate and glutamate via metabolic bypasses. **A.** The parental strains BP265 (*gltAB P_ansAB_-lacZ*) and BP279 (*aspB P_ansAB_-lacZ*) and the strains BP274 (*gltAB citG P_ansAB_-lacZ*), BP276 (*gltAB citG ansR P_ansAB_-lacZ*), and BP292 (*aspB ansR P_ansAB_-lacZ*) were propagated on CGXII plates containing glucose and ammonium as carbon and nitrogen sources, respectively. Aspartate, Asparagine and glutamate were added to a final concentration of 0.5% (w/v). The activity of the *P_ansAB_* promoter was visualized by adding the chromogenic substrate X-Gal to the plates. The plates were incubated for 48 h at 37°C. **B.** Growth of the strains indicated in A. in CGXII liquid medium at 37°C. To remove the amino acids from the precultures, cells were washed twice in 0.9% (w/v) saline solution.

To evaluate the efficiency of ammonium assimilation by the L-aspartase AnsB in the strains BP276 (*gltAB ansR citG*) and BP292 (*aspB ansR*), we cultivated the bacteria in CGXII medium with increasing amounts of ammonium chloride (0 - 302.7 mM). The wild type strain BP264 served as a control. As expected, all strains did not grow in the absence of ammonium (Figure 4A). Albeit forming less biomass than wild type strain, high ammonium concentrations also supported growth of the reconstituted strains BP276 (*gltAB ansR citG*) and BP292 (*aspB ansR*) (Figure 4A). The calculation of the growth rates revealed the assimilation of ammonium via AnsB is limited and less efficient than via the native GS-GOGAT route (Figure 1A and 4B). Moreover, the strain BP276 showed the strongest dependency on ammonium. The lower ammonium dependency of the strain BP292 may be explained by the fact that in the background of the *aspB* aspartate transaminase mutant, the L-aspartase-catalysed ammonium assimilation reaction is only required for *de novo* synthesis of aspartate and asparagine (Figure 1A). In contrast, in the background of *gltAB* GOGAT mutant, the assimilation of ammonium via AnsB is needed for producing glutamate, which is the major amino group donor in the cell. To conclude, albeit less efficient, the alternative entry point for ammonium into central metabolism allows growth of the reconstituted mutants BP276 and BP292.

**Fig. 4.**
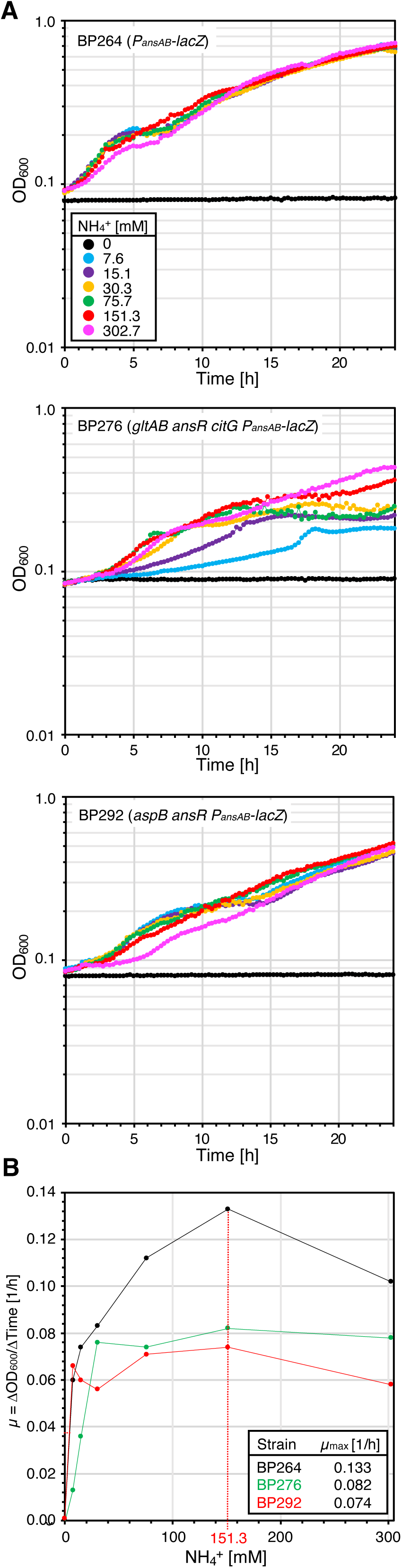
Ammonium dependency of *B. subtilis* strains synthesizing aspartate and glutamate via metabolic bypasses. **A.** The parental strain BP264 (*P_ansAB_-lacZ*) and the strains BP276 (*gltAB ansR citG P_ansAB_-lacZ*) and BP292 (*aspB ansR P_ansAB_-lacZ*) were cultivated at 37°C in SM medium supplemented with increasing amounts of ammonium. **B.** Relationship between the ammonium concentration and the growth rate (µ). The maximum growth rate was reached at an ammonium concentration of 151.3 mM.

As described above, in addition to its role as the major amino group donor in the cell, glutamate serves as the precursor for the synthesis of L-proline, which is as a compatible solute in *B. subtilis* under hyperosmotic growth conditions. Therefore, we assessed the ability of the reconstituted mutants BP276 (*gltAB ansR citG*) and BP292 (*aspB ansR*) to grow under hyperosmotic conditions. For this purpose, the strains including the wild type BP264 as a control were grown in CGXII medium supplemented with increasing amounts of sodium chloride (NaCl). As shown in Figure 5A and 5B, even though the growth of the reconstituted mutants was impaired at elevated NaCl concentrations, both strains synthesize significant amounts of glutamate to withstand an elevated environmental salinity.

**Fig. 5.**
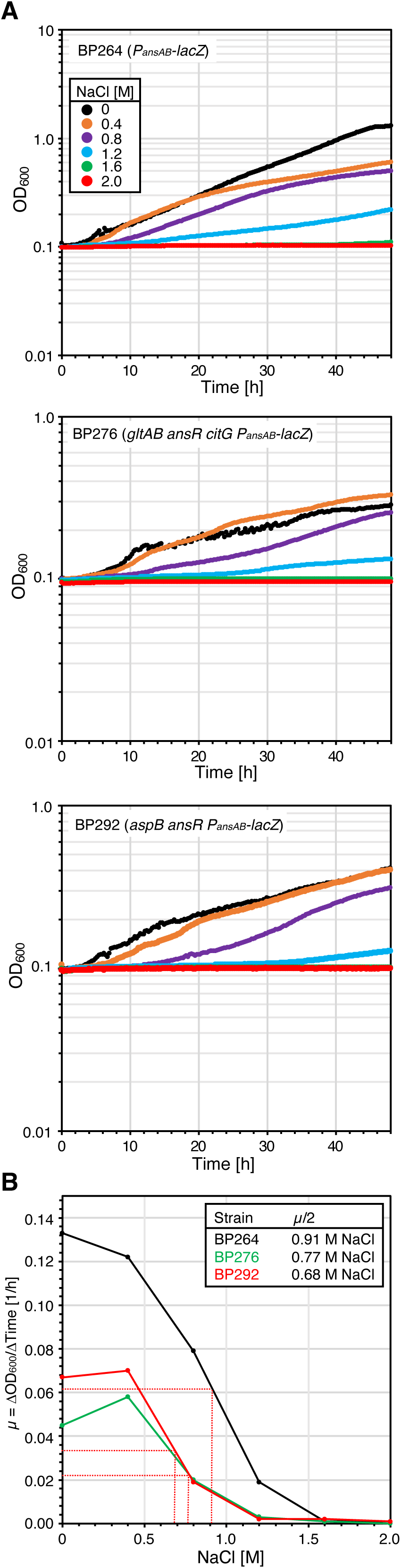
Effect of NaCl on growth of the *B. subtilis* strains synthesizing aspartate and glutamate via metabolic bypasses. **A.** The parental strain BP264 (*P_ansAB_-lacZ*) and the strains BP276 (*gltAB ansR citG P_ansAB_-lacZ*) and BP292 (*aspB ansR P_ansAB_-lacZ*) were cultivated at 37°C in SM medium supplemented with increasing amounts of NaCl. **B.** Relationship between the NaCl concentration and the growth rate (µ).

To determine the relative intracellular concentrations of glutamate, glutamine, aspartate, and asparagine as well as the TCA cycle intermediates citrate, succinate, and malate in the reconstituted mutants BP276 (*gltAB ansR citG*) and BP292 (*aspB ansR*), we performed metabolome analyses. As shown in Figure 6, both strains synthesized similar amounts of glutamate, glutamine, asparagine, and citrate. Moreover, the relative aspartate concentrations were increased and decreased in the strains BP276 and BP292, respectively (Figure 6). It is tempting to speculate that the aspartate concentration is elevated in the strain BP276 because each molecule of glutamate is derived from aspartate. Moreover, due to the deletion of the *citG* gene, the cellular concentration of malate is reduced and allows the strain to direct the metabolic flux from the TCA cycle to the aspartate biosynthetic pathway (Figure 6). In contrast to strain BP276, the presence of the fumarase CitG probably explains a reduced cellular concentration of succinate in the strain BP292. To conclude, despite significant changes in the metabolome of the reconstituted mutants, the metabolic network is robust enough to sustain growth of the bacteria using AnsB for ammonium assimilation.

**Fig. 6.**
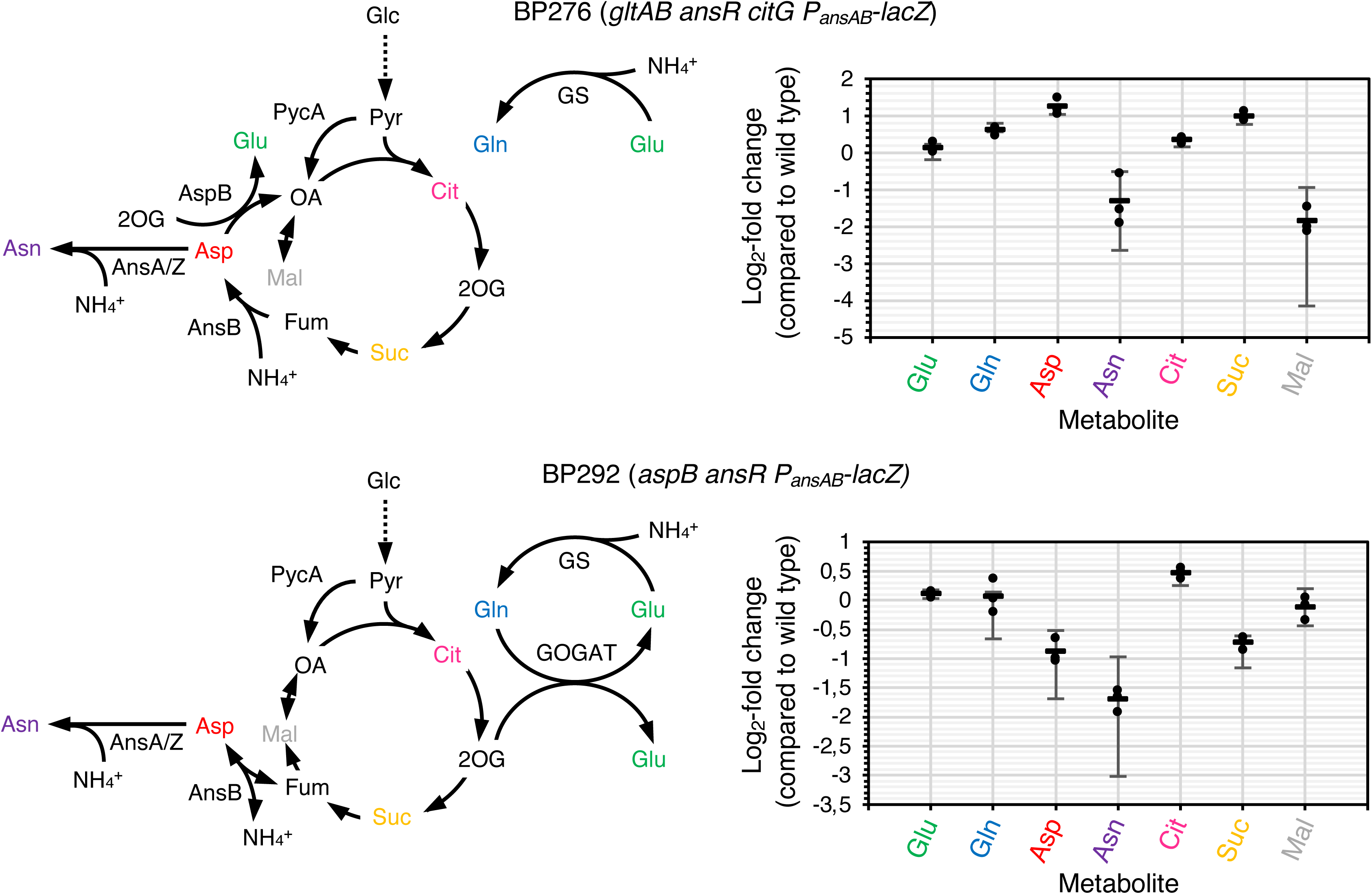
Concentrations of selected metabolites in *B. subtilis* strains synthesizing aspartate and glutamate via metabolic bypasses. The parental strain BP264 (*P_ansAB_-lacZ*) and the strains BP276 (*gltAB ansR citG P_ansAB_-lacZ*) and BP292 (*aspB ansR P_ansAB_-lacZ*) were cultivated at 37°C in SM medium. The metabolites were identified by GC/MS and are shown as log_2_-fold changes compared to the parental strain (see *Experimental procedures*). Mean value and standard deviation of three biological replicates are shown. Glc, glucose; Pyr, pyruvate; Cit, citrate; 2OG, 2-oxoglutarate; Suc, succinate; Fum, fumarate; Mal, malate; OA, oxaloacetate; Asp, aspartate; Asn, asparagine; Glu, glutamate; Gln, glutamine. AnsA and AnsZ, asparaginases; AnsB, aspartase; AspB, aspartate transaminase; GOGAT, glutamate synthase; GS, glutamine synthetase.

### Activation of the *gudB* gene enhances growth of *citG* mutants in rich medium

During the construction of the strains BP275 (*ansR citG P_ansAB_-lacZ*) and BP276 (*gltAB ansR citG P_ansAB_-lacZ*) we observed that the bacteria lacking CitG fumarase activity showed a translucent phenotype on LB rich medium plates. To elucidate the reason for the lytic phenotype, we cultivated the wild type strain BP264 (*P_ansAB_-lacZ*) and the reconstituted mutants BP276 (*gltAB ansR citG P_ansAB_-lacZ*) and BP292 (*aspB ansR P_ansAB_-lacZ*) in LB and in brain heart infusion (BHI) liquid medium. As shown in Figure 7A and 7C, only the strain BP276 lacking the *citG* gene showed a growth defect in LB medium. The fact that the addition of glucose to LB and BHI medium relieves the growth defect of the strain BP276 indicates that the block in the TCA cycle probably prevents the bacteria from efficiently using the amino acids that are present in LB and BHI rich media (Figure 7B and 7D). Next, we performed a short-term evolution experiment by passaging the wild type strain BP264 and the *citG* mutants BP275 (*ansR citG P_ansAB_-lacZ*) and BP276 (*gltAB ansR citG P_ansAB_-lacZ*) for 10 days in LB liquid medium (see *Experimental procedures*). From each culture, we isolated a single colony on LB medium for further analysis. The derivatives of the strains BP264, BP275 and BP276 were designated as BP384, BP369 and BP370, respectively. As shown in Figure 7E, the wild type strain and its evolved derivative were phenotypically indistinguishable from each other. In contrast, the colonies of the evolved *citG* mutants were less translucent than the parental strains (Figure 7E). The cultivation of the bacteria also showed that the lytic phenotype was less pronounced in the *citG* mutants (Figure 7F). Genome sequencing revealed that both evolved *citG* mutant strains had activated the *gudB* GDH gene that is cryptic in the *B. subtilis* laboratory strain SP1 due to a tandem repeat in the open reading frame [Belitsky and Sonenshein, 1998; Zeigler et al., 2003; Gunka et al., 2012]. To conclude, the lytic phenotype of the *citG* mutants can be suppressed by enhancing GDH activity, which is required for efficient utilization of amino acids of the glutamate family.

**Fig. 7.**
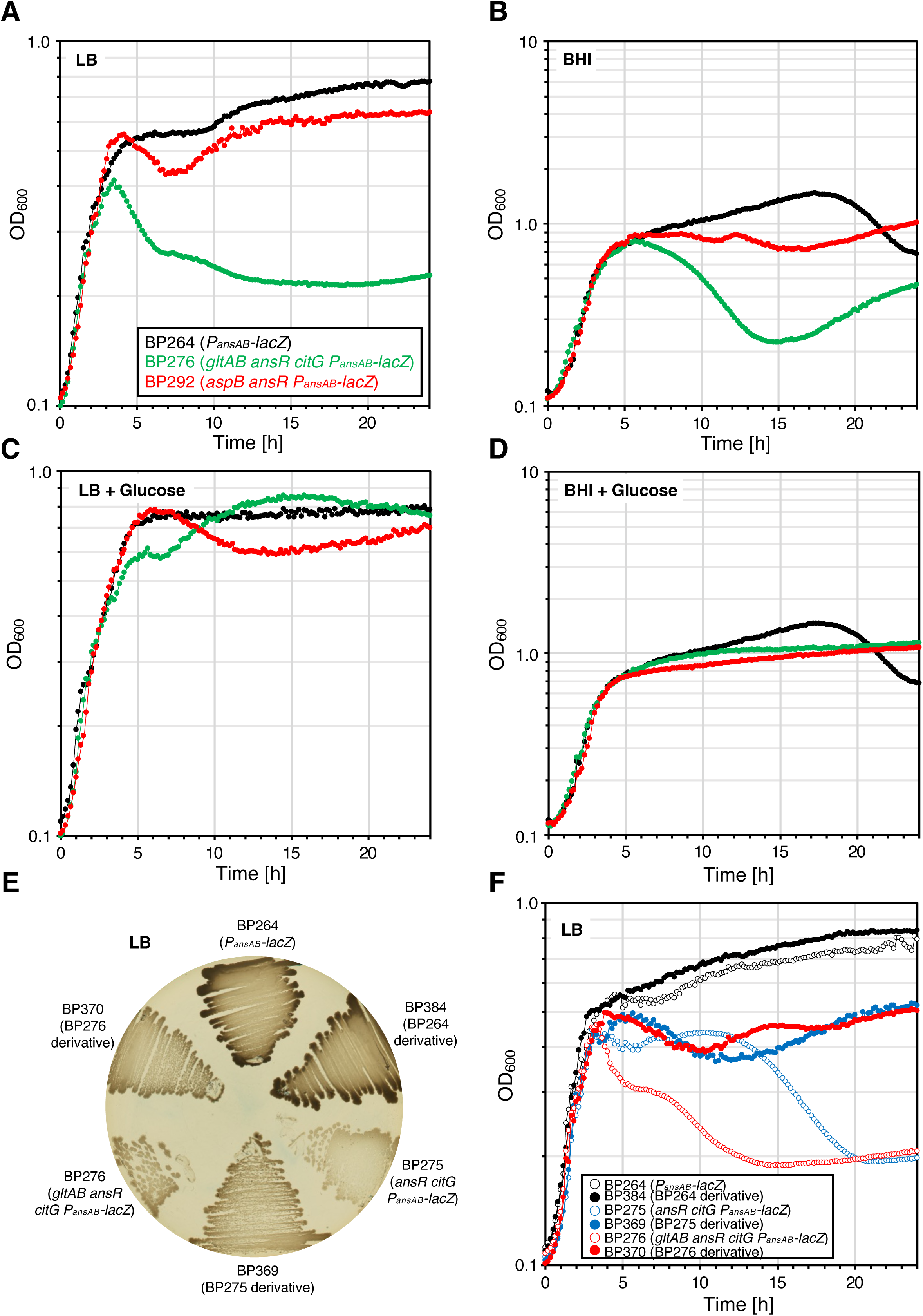
Adaptation of the *B. subtilis* strains synthesizing aspartate and glutamate via metabolic bypasses to rich medium. The parental strain BP264 (*P_ansAB_-lacZ*) and the strains BP276 (*gltAB ansR citG P_ansAB_-lacZ*) and BP292 (*aspB ansR P_ansAB_-lacZ*) were cultivated at 37°C in LB and BHI medium without (**A** and **B**) and with glucose 0.5% (w/v) (**C** and **D**). **E.** Agar plate showing the phenotypes of the strains BP264 (*P_ansAB_-lacZ*), BP276 (*gltAB ansR citG P_ansAB_-lacZ*) and BP292 (*aspB ansR P_ansAB_-lacZ*) and of the strains BP384, BP369 and BP370 that were evolved in LB liquid medium. The plate was incubated for 24 h at 37°C. **F.** Growth of the strains BP264 (*P_ansAB_-lacZ*), BP276 (*gltAB ansR citG P_ansAB_-lacZ*), BP292 (*aspB ansR P_ansAB_-lacZ*), BP384 (LB evolved derivative of BP264), BP369 (LB evolved derivative of BP275) and BP370 (LB evolved derivative of BP276) in LB liquid medium at 37°C.

### Genomic adaptation of *citG* mutants to toxic levels of arginine

The fact that the growth of the *citG* mutants could be improved by the mutational activation of the *gudB* gene suggests that the GDH activity is needed for the utilization of amino acids of the glutamate family and to prevent the accumulation of glutamate to toxic levels. To further address this question, we assessed the toxicity of arginine in the strains lacking CitG fumarase activity. Arginine is taken up by *B. subtilis* via the permeases RocC and RocE and converted to glutamate (Calogero et al., 1994; Gardan et al., 1995, Belitsky and Sonenshein, 1998). For this purpose, we propagated the strains BP264 (*P_ansAB_-lacZ*), BP265 (*gltAB P_ansAB_-lacZ*), BP271 (*ansR P_ansAB_-lacZ*), BP275 (*ansR citG P_ansAB_-lacZ*), BP276 (*gltAB ansR citG P_ansAB_-lacZ*), BP279 (*aspB P_ansAB_-lacZ*), and BP292 (*aspB ansR P_ansAB_-lacZ*) on an LB plate (control) and on SM minimal medium plates supplemented with glucose, glucose plus arginine and only arginine. The strains BP264, BP265, BP271, BP279 and BP292 served as controls. As expected, the strains BP265 and BP271 that are auxotrophic for glutamate and aspartate, respectively, did not grow on the SM plate containing only glucose (Figure 8A). The latter strain also did not grow on the SM plates supplemented with either glucose and arginine or arginine due to the absence of AspB that is required for interconversion of aspartate and glutamate (Figure 1A). Moreover, both *citG* mutants (strains BP275 and BP276) showed the previously observed growth defect on the LB plate and no growth was visible on the SM plate containing only arginine (Figure 8A). Thus, arginine is indeed toxic for the *citG* mutants, probably due to the accumulation of glutamate or TCA cycle intermediates. Moreover, arginine was not toxic for the bacteria in the presence of glucose because the cellular demand for glutamate is higher during growth with the preferred carbon source glucose (Figure 8A) (Belitsky and Sonenshein, 1998; Commichau et al., 2007a). When the arginine-containing plates were further incubated for 48 h at 37°C, we observed that the *citG* mutants BP275 and BP276 formed small and large suppressor mutants, which was not the case for the other strains (Figure 8B; data not shown). Next, we selected two small and two large colonies that were derived from the two *citG* mutants and propagated the bacteria on SM plates supplemented with either glucose and arginine or only arginine (Figure 8C). The wild type strain BP264 and the parental strains BP275 (*ansR citG P_ansAB_-lacZ*) and BP276 (*gltAB ansR citG P_ansAB_-lacZ*) served as controls. As expected, all strains grew on the plates containing glucose that alleviates arginine toxicity. In contrast, the suppressor mutants BP371 - BP374 and BP375 - BP378 that were derived from the strains BP275 and BP276, respectively, showed slight but significant growth on the SM plates supplemented with arginine (Figure 8C). Genome sequencing revealed that the large suppressor mutants BP371, BP372, BP375 and BP376 had acquired mutations that affect the expression of the *aspB* gene (Table 2). In the strains BP371, BP372 and BP376, overproduction of AspB is achieved due to 4-fold amplification of genomic segments that contain the *aspB* gene (Figure 9A). The strain BP375, has acquired a mutation in the *P_dinG_* promoter that lies upstream of the *dinG, ypmA, ypmB* and *aspB* genes. It is tempting to speculate that the mutation in the *P_dinG_* promoter would enhance the expression of *dinG* including the *aspB* gene. Since AspB can convert glutamate and oxaloacetate to aspartate and 2-oxoglutarate, the overproduction of the aspartate transaminase would prevent the accumulation of glutamate to toxic levels. Previously, it has also been observed that *a B. subtilis* mutant lacking GDH activity and AnsR can grow with glutamate as the sole source of carbon and nitrogen [Flórez et al., 2010]. The two other suppressors BP373 and BP374 that were derived from the strain BP275 (*ansR citG P_ansAB_-lacZ*) and formed small colonies on arginine-containing SM plates had accumulated mutations in the *rocC* gene encoding the arginine permease RocC (Figure 1A) (Table 2). The G1232A nucleotide exchange in the *rocC* allele of the strain BP373 causes the P411L exchange in RocC. Since the residue P411 is located in the transmembrane helix 11 in RocC, it is very likely that P411L exchange reduces the activity of the arginine permease. The insertion of T at the position 698 in the suppressor BP374 results in a frameshift and inactivates RocC due to a truncation by 270 amino acids. Thus, in addition to *aspB* overexpression, the bacteria lacking CitG activity can adapt to toxic levels of glutamate by reducing arginine uptake. It remains elusive why BP373 carries a mutation in the *acpP* gene encoding the essential AcpA protein that is involved in fatty acid biosynthesis (Table 2) [Schujman et al., 2003]. It is tempting to speculate that the D19Y replacement in AcpA could affect the interaction between AcpA and the acyl-carrier protein synthase AcpS. Moreover, Genome sequencing identified mutations in the *P_odhA_* promoter region and in the *odhA* gene in the strains BP377 and BP378, respectively, that were derived from the strain BP276 (*gltAB ansR citG P_ansAB_-lacZ*) (Table 2). Since the deletion in the *odhA* gene of the strain BP378 certainly inactivates the OdhAB-PdhD 2-oxoglutarate dehydrogenase, it is likely that the mutation in the *P_odhA_* promoter region reduces the expression of the *odhA* gene. The strains BP377 and BP378 also carry mutations in the essential *pyrH* gene encoding the uridylate kinase PyrH that converts ATP and UMP to ADP and UDP. Since the residue T145 is in α-helix 4 in PyrH, the T145P replacement certainly affects the activity of the enzyme and thus probably the cellular concentration of the nucleotides (Table 2) [Quinn et al., 1991]. To conclude, arginine toxicity can be relieved by AspB overexpression, by reducing arginine uptake and by decreasing the metabolic flux through the TCA cycle.

**Fig. 8.**
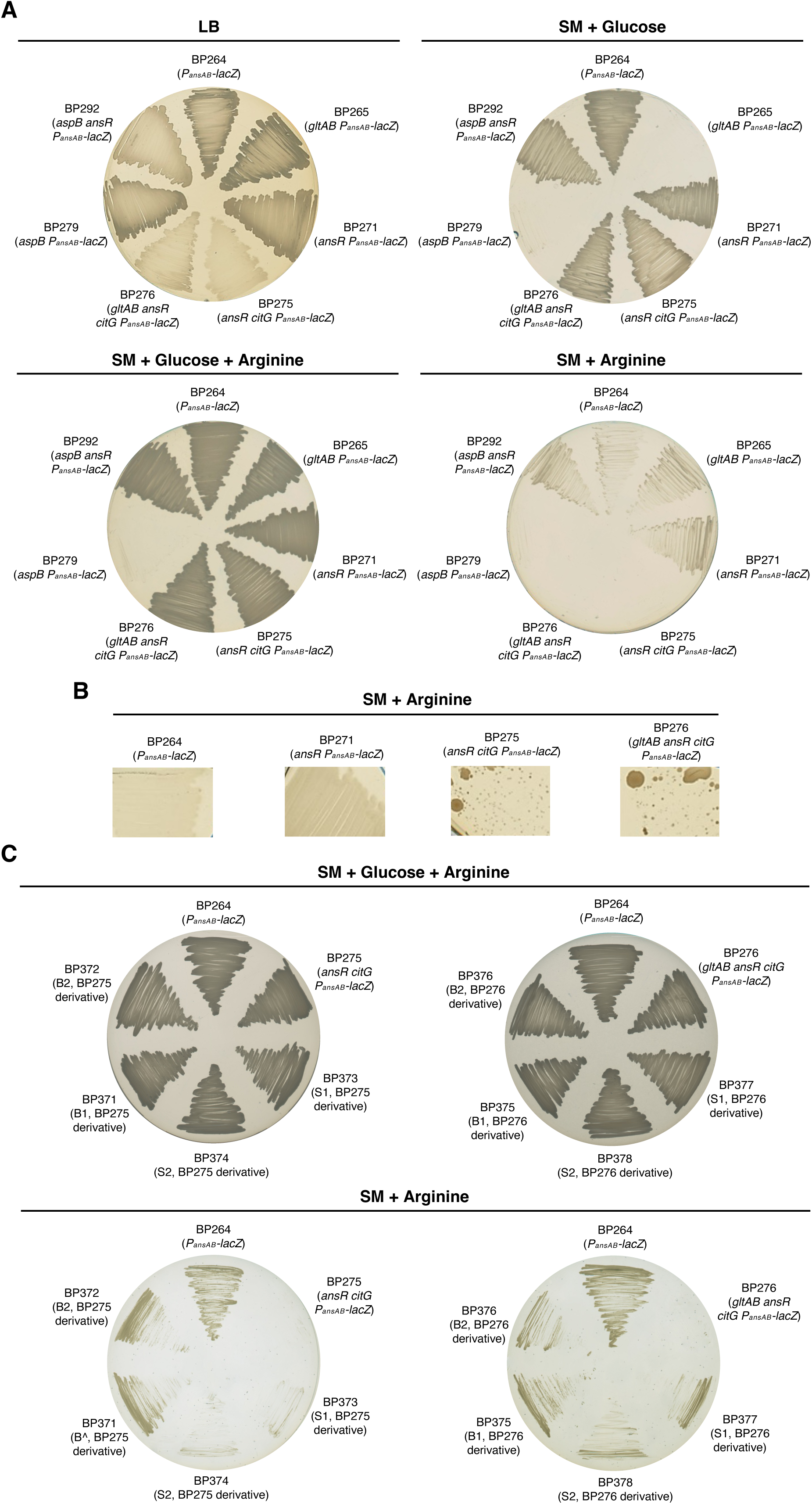
Adaptation of *B. subtilis* fumarase mutants to arginine. **A.** Growth of the strains BP264 (*P_ansAB_*-*lacZ*), BP265 (*gltAB P_ansAB_*-*lacZ*), BP271 (*ansR P_ansAB_*-*lacZ*), BP275 (*ansR citG P_ansAB_*-*lacZ*), BP276 (*gltAB ansR citG P_ansAB_*-*lacZ*), BP279 (*aspB P_ansAB_*-*lacZ*), and BP292 (*aspB ansR P_ansAB_*-*lacZ*) on LB and SM plates. Arginine and glucose were added to a final concentration of 0.5% (w/v). **B.** Emergence of suppressor mutants derived from the fumarase mutants BP275 (*ansR citG P_ansAB_*-*lacZ*) and BP276 (*gltAB ansR citG P_ansAB_*-*lacZ*) on SM plates containing 0.5% (w/v) arginine. The plates were incubated for 10 days at 37°C. **C.** Growth of the arginine adapted suppressor mutants BP371 (B1, BP275 derivative), BP372 (B2, BP275 derivative), BP373 (S1, BP275 derivative), and BP374 (S1, BP275 derivative), BP375 (B1, BP276 derivative), BP376 (B2, BP276 derivative), BP377 (S1, BP276 derivative), and BP378 (S2, BP276 derivative) on SM-glucose plates without and with 0.5% (w/v) arginine. The strains BP264 (*P_ansAB_*-*lacZ*), BP275 (*ansR citG P_ansAB_*-*lacZ*), and BP276 (*gltAB ansR citG P_ansAB_*-*lacZ*) served as controls. The plates were incubated for 48 h at 37°C. “S” and “B” indicate “small colony morphology” and “big colony morphology”, respectively.

**Fig. 9.**
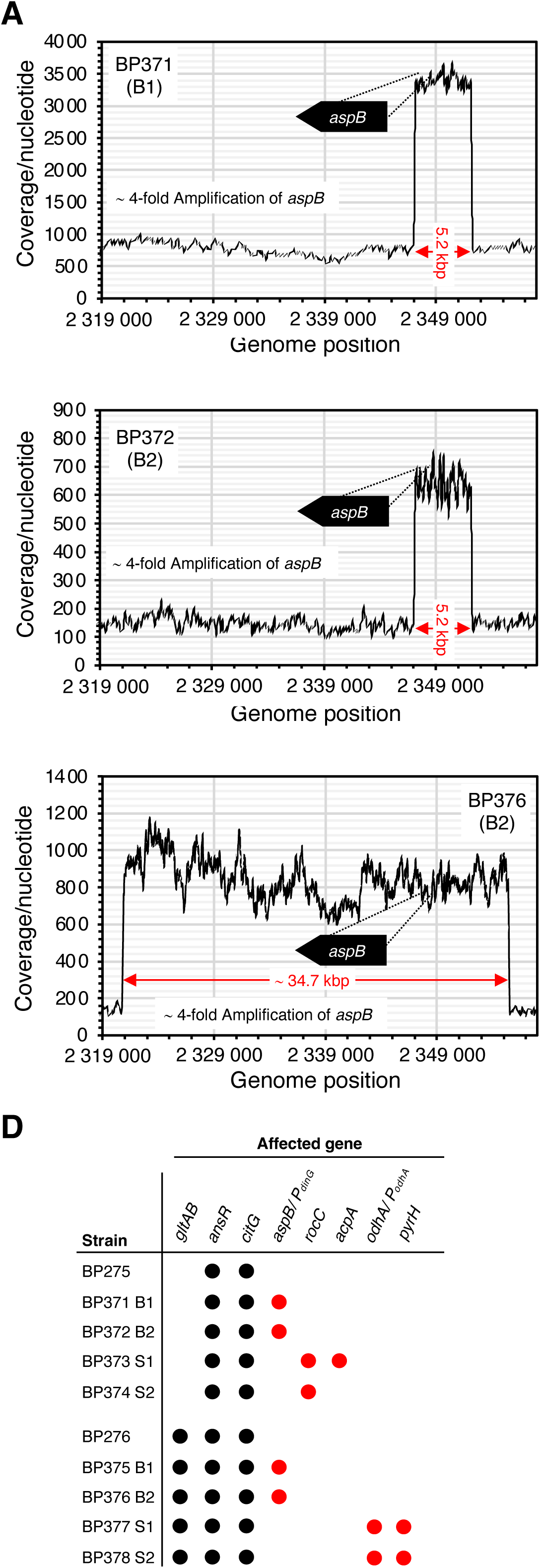
Genomic adaptation of *B. subtilis* fumarase mutants relieves arginine toxicity. **A.** Read coverages along the chromosomal segment ranging from 2.319.000 to 2.358.000 bp. Based on the average coverage of the amplified regions and of the remaining genomes it can be inferred that four copies of the 5.2 and 37.4 kbp long regions containing the *aspB* gene are present in the suppressors BP371 (B1, BP275 derivative), BP372 (B2, BP275 derivative), and BP376 (B2, BP276 derivative). **B.** Genes that are absent (black circles) or mutated (red circles) in the arginine adapted suppressor mutants BP371 (B1, BP275 derivative), BP372 (B2, BP275 derivative), BP373 (S1, BP275 derivative), and BP374 (S1, BP275 derivative), BP375 (B1, BP276 derivative), BP376 (B2, BP276 derivative), BP377 (S1, BP276 derivative), and BP378 (S2, BP276 derivative).

**Table 2:** Identified mutations in the evolved *ansR citG* and *ansR citG gltAB* mutants.

|  |  |  |  |  |  |
| --- | --- | --- | --- | --- | --- |
| **Mutations in the strains BP275 (*ansR citG P<sub>ansAB-lacZ</sub>*) and BP276 (*gltAB ansR citG P<sub>ansAB-lacZ</sub>*) evolved in LB medium** |  |  |  |  |  |
| **Strain** | **Mutant** | **Parental strain** | **Phenotype** | **Affected gene, mutations** | **Amino acid exchanges, effect on the protein** |
| BP384 | LBW<sup>A</sup> | BP264 (*P<sub>ansAB-lacZ</sub>*) | Growth in LB | - | - |
| BP369 | LB1<sup>A</sup> | BP275 (*ansR citG P<sub>ansAB-lacZ</sub>*) | Growth in LB | *gudB* ΔG279-C287 | Synthesis of GudB1 |
| BP370 | LB2<sup>A</sup> | BP276 (*gltAB ansR citG P<sub>ansAB-lacZ</sub>*) | Growth in LB | *gudB* ΔG279-C287 | Synthesis of GudB1 |
| **Mutations reducing arginine toxicity of the strains BP275 (*ansR citG P<sub>ansAB-lacZ</sub>*) and BP276 (*gltAB ansR citG P<sub>ansAB-lacZ</sub>*)** |  |  |  |  |  |
| BP371 | B1<sup>A</sup> | BP275 (*ansR citG P<sub>ansAB-lacZ</sub>*) | Growth on SM-Arg, big colonies | 5.2 kbp amplification including *aspB* | Increased AnsB synthesis |
| BP372 | B2<sup>A</sup> | BP275 (*ansR citG P<sub>ansAB-lacZ</sub>*) | Growth on SM-Arg, big colonies | 5.2 kbp amplification including *aspB* | Increased AnsB synthesis |
| BP373 | S1<sup>A</sup> | BP275 (*ansR citG P<sub>ansAB-lacZ</sub>*) | Growth on SM-Arg, small colonies | *acpA* G55T *rocC* C1232T | AcpA D19Y, RocC P411L |
| BP374 | S2<sup>A</sup> | BP275 (*ansR citG P<sub>ansAB-lacZ</sub>*) | Growth on SM-Arg, small colonies | *rocC* +T698 | RocC ΔC270 |
| BP375 | B1<sup>A</sup> | BP276 (*gltAB ansR citG P<sub>ansAB-lacZ</sub>*) | Growth on SM-Arg, big colonies | *P<sub>dinG</sub>* (C-63T) | Increased AspB synthesis |
| BP376 | B2<sup>A</sup> | BP276 (*gltAB ansR citG P<sub>ansAB-lacZ</sub>*) | Growth on SM-Arg, big colonies | 34.7 kbp amplification including *aspB* | Increased AspB synthesis |
| BP377 | S1<sup>A</sup> | BP276 (*gltAB ansR citG P<sub>ansAB-lacZ</sub>*) | Growth on SM-Arg, small colonies | *odhA* ΔT175-A351, *pyrH* A433C | OdhA not synthesized, PyrH T145P |
| BP378 | S2<sup>A</sup> | BP276 (*gltAB ansR citG P<sub>ansAB-lacZ</sub>*) | Growth on SM-Arg, small colonies | *P<sub>odhA</sub>* (G-189A), *pyrH* A433C | Altered *odhA* expression? PyrH T145P |

1021 <sup>A</sup>Mutations were identified by genome sequencing.

## Discussion

Here, we found that the glutamate auxotrophy of a *B. subtilis gltAB* mutant lacking GOGAT activity can be relieved by the mutational inactivation of the *ansR* and *citG* genes. The de-repression of the *ansB* gene and the disruption of the TCA cycle allows the *gltAB* mutant to synthesize glutamate via the L-aspartase/L-aspartate transaminase-dependent route (Figures 1A and 1D). We also show that the inactivation of the *ansR* gene is sufficient to restore the aspartate prototrophy of an *ansB* mutant. It is tempting to speculate that the lack of AnsR is sufficient to allow growth of the *aspB* mutant because the cell needs significantly less aspartate than glutamate. This hypothesis is underlined by the fact that glutamate is indeed the dominant cellular metabolite in prokaryotes and in eukaryotes [Bennett et al., 2009].

Previously, it has been shown that also other genetic lesions can lead to glutamate auxotrophy in *B. subtilis*. For instance, the inactivation of the *gltC* gene encoding the DNA-binding regulator GltC that is required for the transcriptional activation of the *gltAB* genes results in glutamate auxotrophy (Figure 1B) [Bohannon and Sonenshein, 1989; Dormeyer et al., 2017]. Interestingly, the expression of the *gltAB* genes can be quickly restored by (i) a gain-of-function mutation in the *gltR* gene, encoding a LysR-type transcription factor of unknown function [Belitsky and Sonenshein, 1997; Dormeyer et al., 2017], (ii) by a promoter-up mutation in the *P_gltAB_* promoter, and (iii) by the selective amplification of a chromosomal segment containing the *gltAB* genes [Dormeyer et al., 2017]. It has also been shown that a *B. subtilis ccpA* mutant is auxotrophic for glutamate due to insufficient expression of the *gltAB* genes and de-repression of the *rocG* GDH gene that is repressed by CcpA during growth with glucose [Faires et al., 1999; Wacker et al., 2003, Commichau et al., 2007a;]. Strikingly, even in the absence of the pleiotropic transcription factor CcpA, *B. subtilis* may restore glutamate biosynthesis by the accumulation of mutations in the *topA* gene encoding the essential DNA topoisomerase I TopA [Reuß et al., 2018]. The TopA mutant variants cause glutamate prototrophy of the *ccpA* mutant due to enhanced relaxation of the chromosomal DNA, which results in the re-organization of the global transcription network, re-routing of central metabolism and in the inactivation of the GDH [Reuß et al., 2018]. Thus, the glutamate auxotrophy in *B. subtilis* can be relieved by mutations restoring the expression of the *gltAB* genes or by the mutational activation of an alternative glutamate biosynthesis route. It has also been demonstrated that the function of an interrupted metabolic pathway can be restored by the overexpression of a native enzyme that is usually involved in another unrelated metabolic pathway. In *B. subtilis*, the γ-glutamyl-phosphate reductase ProA is involved in the biosynthesis proline that serves as a building block and a compatible solute during osmotic stress [Bremer and Krämer, 2019]. Thus, the growth of a *proA* mutant should be strictly proline dependent. However, the *proA* mutant rapidly forms suppressor mutants that overexpress the ornithine aminotransferase RocD that shares no sequence similarity with ProA but synthesizes the same reaction product [Zaprasis et al., 2014; Stecker et al., 2022]. Recently, it has been observed that pre-existing enzymatic activities can bypass genetic lesions that block the metabolic flux via the Embden-Meyerhof-Parnas pathway in *E. coli* [Iacometti et al., 2022]. These metabolic bypasses may proceed either via methylglyoxal, a highly toxic intermediate, or via enzymes involved in serine metabolism [Iacometti et al., 2022]. Thus, in phylogenetically distantly related organisms, enzymes that are usually involved in different metabolic pathways can be recruited to rewire the metabolic network.

There are several other options to bypass disrupted metabolic pathways in bacteria [Kim and Copley, 2007]. For instance, an enzymatic reaction that is absent due to a gene deletion can be circumvented by recruiting a promiscuous enzyme that catalyses the missing reaction. Indeed, in *B. subtilis* it has been observed that the overexpression of the threonine synthase ThrC allows growth of an *ilvA* mutant that lacks the major threonine dehydratase IlvA [Rosenberg et al., 2016]. Due to sequence homology and similar catalytic mechanisms, it has been suggested that threonine dehydratase and threonine synthase evolved from a common ancestor enzyme [Parsot, 1986]. However, since ThrC has a minor threonine dehydratase activity, a *B. subtilis ilvA* mutant overexpressing the *thrC* gene only grows in the presence of threonine. A missing enzymatic reaction can also be circumvented by promiscuous enzymes that divert intermediates from other metabolic pathways to products, which feed into the disrupted pathway [Kim et al., 2010, 2019; Cotton et al., 2020]. In *E. coli* it has been shown that a 4-step bypass pathway, which is patched together from promiscuous enzymes relieves vitamin B6 auxotrophy of a *pdxB* mutant lacking the erythronate 4-phosphate dehydrogenase PdxB [Kim et al., 2010, 2019]. The flexibility of the cellular metabolic network even allows the evolution of a non-native and truncated biosynthetic pathway. For instance, in *B. subtilis*, two genomic alterations are sufficient to enable growth of a vitamin B6 auxotrophic *B. subtilis* mutant synthesizing the enzymes that catalyse the last two steps of the vitamin B6 biosynthesis pathway in *E. coli* [Rosenberg et al., 2018; Rosenberg and Commichau, 2019; Richts and Commichau, 2021]. Finally, metabolic systems can overcome pathway damage by extensively rerouting metabolic pathways and modifying existing enzymes for unnatural functions [Pontrelli et al., 2018; Veeravalli et al., 2010].

The present work also raises the question of whether the GS-GOGAT-dependent biosynthesis pathway is the dominant pathway for the biosynthesis of the major cellular amino group donor in nature because the non-canonical fumarate-based ammonium assimilation pathway does not depend on ATP and NADPH_2_ (Figure 1D). Like in *B. subtilis*, also in *E. coli* the overexpression of the native L-aspartase can replace the canonical glutamate-based ammonium assimilation pathway [Schulz-Mirbach et al., 2022]. To address the question of how widespread the coding capacity for the bypass is, we performed a bioinformatic survey. We used the GltAB, AnsB, AspB and CitG proteins from *B. subtilis* as queries to screen all annotated proteins from 14,954 bacterial genomes. This screen revealed that > 1600 genomes lack homologs for GltA and GltB but possess AsnB, AspB and CitG homologs (Figure S3). It will be interesting to test whether the L-aspartase/L-aspartate transaminase-dependent route is the major glutamate biosynthesis pathway in these species. Previously, it has been reported that a GOGAT/GDH-deficient *Corynebacterium glutamicum* strain still grows with glucose and ammonium as single sources of carbon and nitrogen, respectively (Figure S4) [Rehm et al., 2010]. However, the Aspartase AspA, which shares 46.7% overall sequence identity with AnsB from *B. subtilis*, is not involved in the fumarate-based ammonium assimilation pathway because a *C. glutamicum* mutant lacking GOGAT and GDH activity as well as AspA still grows on minimial medium plates containing only glucose and ammonium (Figure S4). This suggests the existence of another, yet unknown route for glutamate biosynthesis in this organism.

As described above, glutamate biosynthesis and degradation are tightly regulated depending on the carbon and nitrogen sources that are available for *B. subtilis* (Figure 1) [Gunka and Commichau, 2012]. Here we also observed that rich medium and the presence of arginine inhibits growth of the *B. subtilis* strain relying on the L-aspartase/L-aspartate transaminase-dependent route for glutamate biosynthesis. The growth defect can be relieved by increased synthesis of AspB, by reduced flux through the TCA cycle and arginine uptake. In the future it will be interesting to investigate whether the *B. subtilis* strain using the alternative glutamate biosynthesis route can be evolved in such a way that it robustly grows during nitrogen limitation and excess.

### Experimental procedures

#### Bacterial strains, chemicals, and DNA manipulation

Bacterial strains used in this study are listed in Table S1. Primers were purchased from Sigma-Aldrich (Munich, Germany) and are listed in Table S2. Chemicals and media were purchased from Sigma-Aldrich (Munich, Germany), Carl Roth (Karlsruhe, Germany) and Becton Dickinson (Heidelberg, Germany). Bacterial chromosomal DNA was isolated using the peqGOLD bacterial DNA kit (Peqlab, Erlangen, Germany). PCR products were purified using the PCR purification kit (Qiagen, Germany). Phusion DNA polymerase was purchased from Thermo Scientific (Germany) and used according to the manufacturer’s instructions.

#### Cultivation of bacteria

Bacteria were grown in lysogeny broth (LB) [Sezonov et al., 2007] and brain heart infusion (BHI) [Rosenow, 1919] rich medium or in Spizizen (SM) [Anagnostopoulos and Spizizen, 1961], MSSM [Gundlach et al., 2017], C-Glc [Commichau et al., 2007; Dormeyer et al., 2019] and CGXII [Keilhauer et al., 1993] minimal medium. Agar plates were prepared with 15 g agar/l (Roth, Germany). Growth in liquid medium was monitored using 96-well plates (Microtest Plate 96-Well, F Sarstedt, Germany) at 37°C and medium orbital shaking at 237 cpm (4 mm) in a Synergy H1 plate reader (Agilent, USA) equipped with the Gen5 software, and the OD_600_ was measured in 10 - 15 min intervals. Single colonies were used to inoculate 5 ml overnight LB cultures that were incubated at 220 rpm and 30°C. The OD_600_ was adjusted to 0.1 and 150 µl of the cell suspensions were transferred into 96-well plates. Bacteria were cultivated in the Synergy H1 plate reader as described above.

#### Plasmid and strain construction

The plasmids used and generated in this study are listed in Table S3. The plasmids pBP1110 and pBP1111 carrying the carrying the *P_ansAB_* promoter and the *ansR* gene together with the *P_ansAB_* promoter, respectively, were constructed as follows. The *P_ansAB_* promoter and the *ansR-P_ansAB_* promoter fragments were amplified by PCR using the primer pairs SM1/SM2 and SM36/SM2, respectively, digested with *Eco*RI and *Bam*HI and ligated to pAC7 [Weinrauch et al., 1991] that was cut with the same enzymes. The generated plasmids were verified by Sanger sequencing (Microsynth-SeqLab Sequence Laboratories).

Deletion of the *ansAB, ansR, aspB, citG, gltAB*, and *recN* genes in *B. subtilis* was achieved by transformation with long-flanking homology (LFH) PCR products constructed using oligonucleotides (Table S2) to amplify DNA fragments flanking the target gene and the intervening *aad3* spectinomycin, *cat* chloramphenicol, and *ermC* erythromycin/lincomycin resistance genes from the plasmids pDG1726, pGEM-cat, and pDG647 [Guérout-Fleury et al., 1995], respectively, as described previously [Gaballa et al., 2010]. When required, antibiotics were added to the following concentrations: ampicillin (100 µg/ml), kanamycin (10 µg/ml), chloramphenicol (5 µg/ml), spectinomycin (150 µg/ml), erythromycin and lincomycin (2 µg/ml and 25 µg/ml). *B. subtilis* was transformed with plasmids, PCR products and with chromosomal DNA according to a previously described two-step protocol [Kunst and Rapoport, 1995]. AmyE amylase activity was detected after growth on agar plates containing nutrient broth (7.5 g/l) Bacto agar (17 g/l) (Difco) and hydrolysed starch (5 g/l) (Connaught). Starch degradation was detected by sublimating iodine onto the plates.

#### Genome sequencing

Genomic DNA was prepared from 500 µl overnight cultures using the MasterPure Complete DNA & RNA Purification Kit (Lucigen, Middleton, USA) following the instruction of the manufacturer with the modification of physically opening cells with the TissueLyser II (Qiagen). Purified genomic DNA was paired end sequenced (2 x 150 bp) (GENEWIZ). The reads were mapped onto the *B. subtilis* reference genome NC_000964 from GenBank [Barbe et al., 2009] as previously described [Widderich et al., 2016] using the Geneious software package (Biomatters Ltd.) [Kearse et al., 2012]. All identified mutations were verified by performing PCRs and Sanger sequencing.

#### Metabolomics

The *B. subtilis* strains were grown overnight in 4 ml LB medium at 28°C and 160 rpm. The overnight cultures were used to inoculate 100 ml shake flasks containing 10 ml SM medium to an OD_600_ of 0.1. The cultures were incubated at 37°C and 160 rpm. 0.5 mg biomass were collected using a PVDF filter (0.45 µm pore site) in a glass frit. The cells on the filters were resuspended in 1 ml ice-cold extraction solution (acetonitrile/methanol/ultrapure water, 40%/40%/20%) and incubated for 1 h at -20°C. The cell extracts were centrifuged for 15 min and 20,000 g at - 9°C, and stored at – 80°C until further processing. Relative concentrations of the metabolites Glu, Gln, Asp, Asn, Cit, Suc and Mal in the cell extracts were measured via isotope ratio LC-MS/MS as described previously [Guder et al. 2017].

#### Evolution of B. subtilis in LB medium

100 ml shake flasks containing 10 ml LB medium were inoculated with a single colonies of the *B. subtilis* strains BP264, BP275 and BP276 and cultivated for 24 h at 37°C. Next day, 100 µl of the cultures were used to inoculate a fresh shake flask. This step was repeated 10 times. Single colonies were isolated by propagating the aliquots of the cultures on LB plates. After phenotypic inspection, one derivative of each strain (Figure 7E) was subjected to genome sequencing.

#### Identification of bacterial genomes lacking the B. subtilis GOGAT

Annotated proteins from all available bacterial genomes were downloaded from RefSeq (14,954 genomes as of 13.04.2022). We used BLASTP searches to identify genomes lacking GltAB but possessing AnsB, AspB and CitG. In practice, bacterial genomes with the *B. subtilis* GltA and GltB subunits were identified, and the enzyme was considered to be absent if none of the subunits had a significant BLASTP hit. *B. subtilis* proteins were used as queries and an e-value threshold of e-50 was applied, experimentally tested on several bacterial species. 2791 genomes were found to lack GltAB, which were subjected to further BLASTP searches to identify AnsB, AspB and CitG using the *B. subtilis* protein queries (e-value<e-50). 1642 genomes lacked GtlAB but had AnsB, AspB and CitG homologs. The taxonomic lineage of these genomes was obtained from NCBI through the use of taxIDs in ETE3 [Huerta-Cepas et al. 2016].

## Acknowledgements

We are grateful to the members of the Commichau laboratory for fruitful comments and suggestions. We thank Janina Berg, Renato Carrillo and Karin Krumbach for the help with some experiments. This project has received funding from the DFG (Co 1139/3-1 to F.M.C.).

## Conflict of interest

The authors declare that there is no potential conflict of interest.

## Supporting information

Additional support information may be found in the online version of the article at the publisher’s web-site:

---

**Fig. S1.**
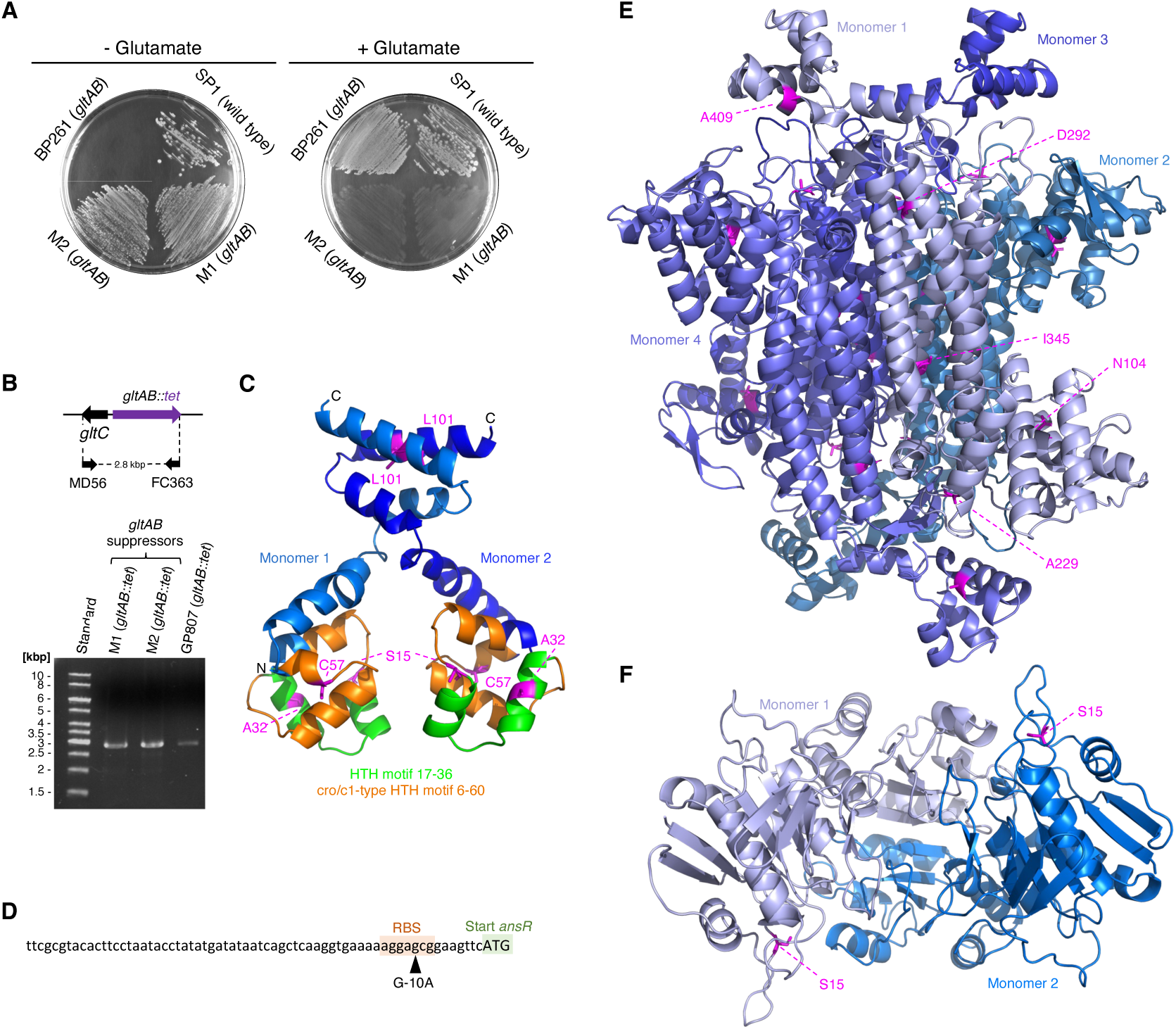
**A.** Growth of the *gltAB* suppressor mutants BP364 (M1) and BP365 (M2), and the parental strain BP261 (*gltAB*) on CGXII plates without and with 0.5% (w/v) glutamate. The plates were incubated for 48 h at 37°C. **B.** Verification of the replacement of the *gltAB* genes by the *tet* resistance gene in the suppressors BP364 (M1) and BP365 (M2). Chromosomal DNA of the *gltAB* mutant GP807 served as the control. **C.** Localization of amino acid exchanges in a structure model of AnsR. The model was generated using the Swiss-model server for homology modeling of protein structures [Waterhouse et al., 2018] and a model of the EspR transcription factor from *Mycobacterium tuberculosis* (PDBid: 3QF3) [Blasco et al., 2011]. **D.** Mutation in the ribosome binding site (RBS) of the *P_ansR_* promoter in the suppressor mutant BP287 that was derived from the strain BP265 (*gltAB P_ansAB_-lacZ*). **E** and **F**, localization of amino acid exchanges in a structure model of CitG and AnsA, respectively. The models were generated as described for the AnsR model using structures of the *E. coli* fumarase (PDBid: 6P3C) and the *Thermococcus kodakarensis* L-asparaginase (PDBid: 5Ot0) [Guo et al., 2017].

**Fig. S2.**
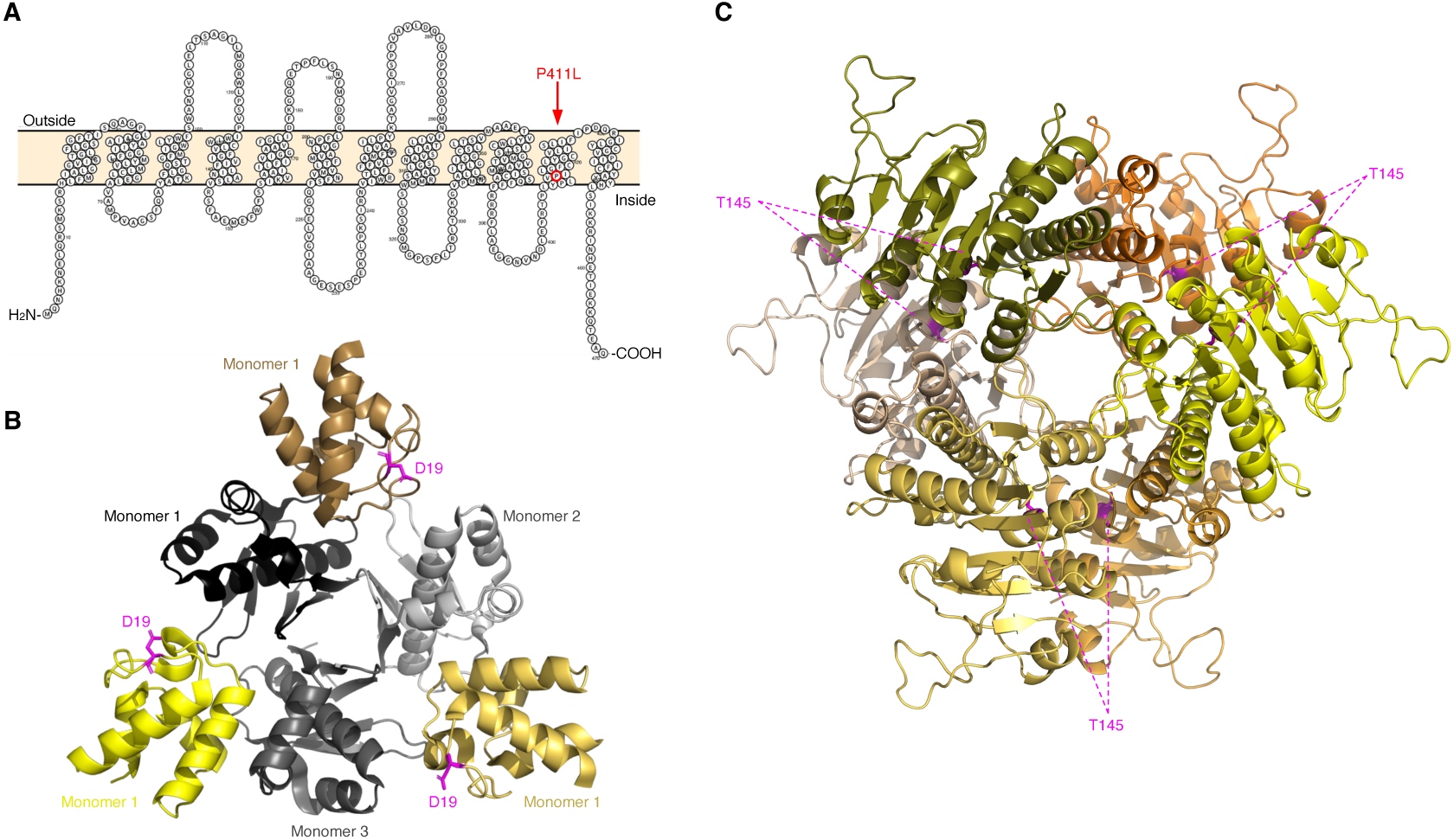
**A.** Localization of the P411L exchange in RocC of the suppressor mutant BP373. The RocC topology model was created with Protter [Omasists et al., 2014]. **B** and **C**, localization of amino acid exchanges in a structure model of the AcpA-AcpS complex and of PyrH, respectively. The models were generated as described for the AnsR model using structures of *B. subtilis* AcpA-AcpS (PDBid: 1F80) [Parris et al., 2000] and the *E. coli* UMP kinase (PDBid: 2V4Y) [Meyer et al., 2008].

**Fig. S3.**
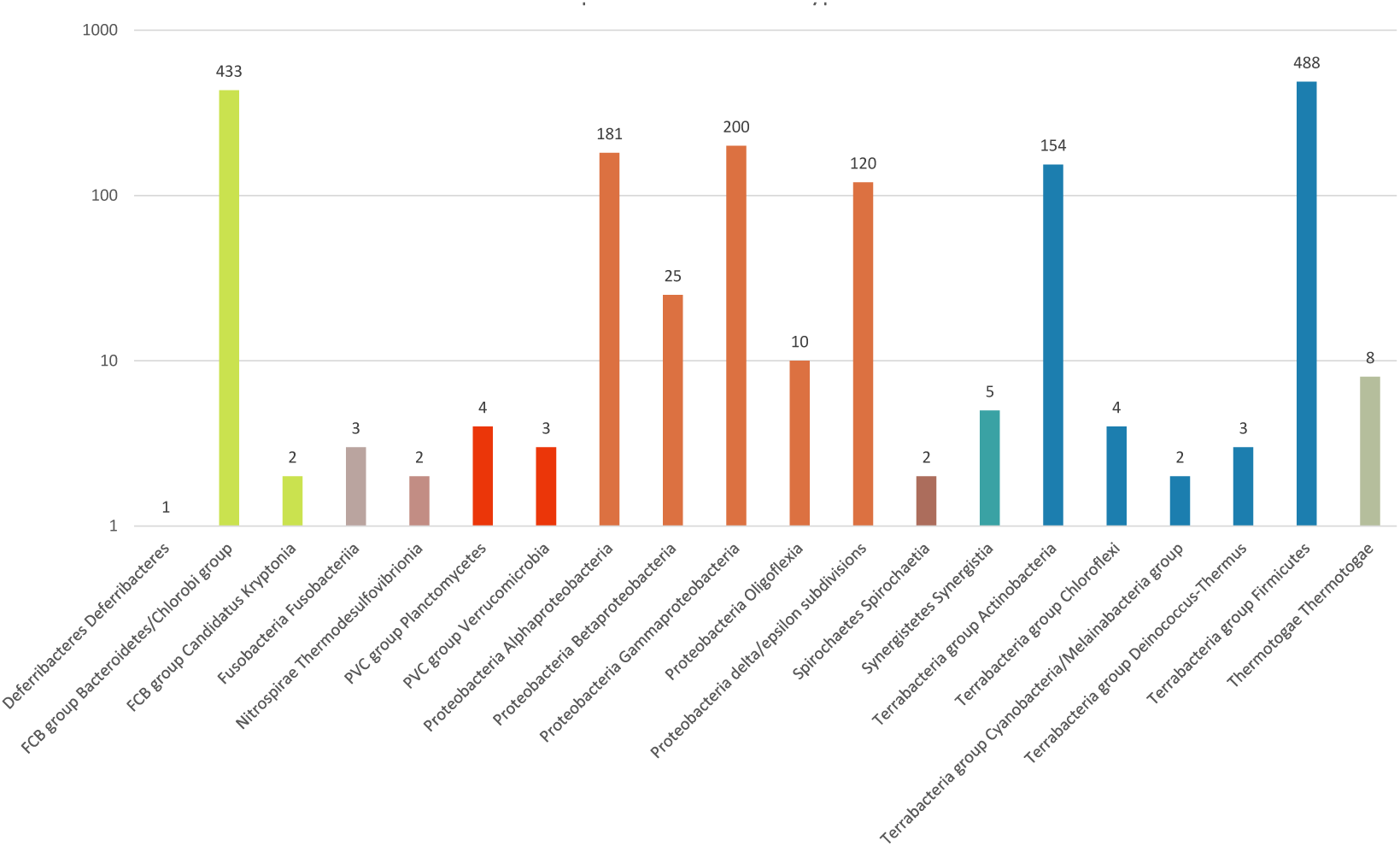
Taxonomic distribution of bacterial species lacking GltA and GltB homologs and possessing AnsB, AspB and CitG homologs, thus suggesting a possible metabolic bypass (in a total of 1642 from 14954 investigated genomes).

**Fig. S4.**
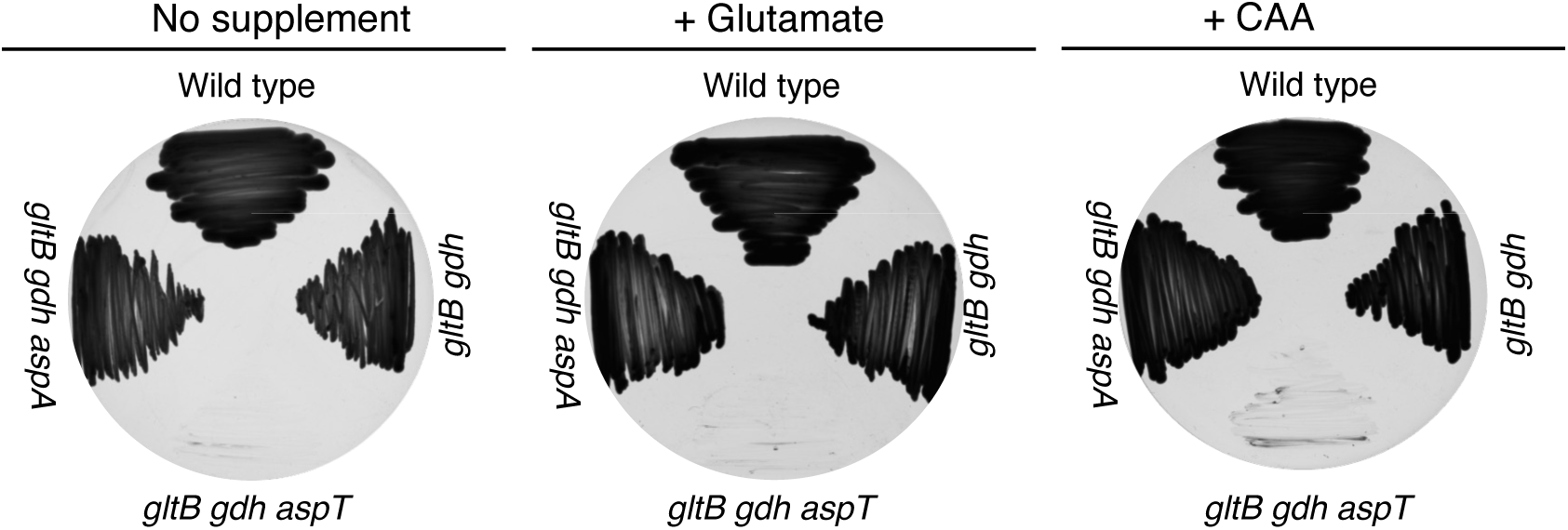
Growth of the *C. glutamicum* wild type and the *gltB gdh*, *gltB gdh aspT* and *gltB gdh aspA* mutants on CGXII medium in the absence and in the presence of glutamate (0.5% (w/v)) or casamino acid hydrolysate (CAA, 0.1% (w/v)). The plates were incubated for 48 h at 30°C. The *C. glutamicum* aminotransferases AspT and AspA share 24% and 46.7% overall amino acid sequence identity, respectively, with the *B. subtilis* AspB and AnsB proteins respectively.

**Table S1.** Strains.

| Strain | Bacterium, mutant | Genotype | Reference, construction<sup>a</sup> |
| --- | --- | --- | --- |
| SP1 | *B. subtilis* | Prototrophic derivative of strain 168 | Richts et al., 2020 |
| 168 | *B. subtilis* | *trpC2* | Laboratory strain collection |
| GP807 | *B. subtilis* | *trpC2 gltAB::tet* | LFH → 168 |
| GP1153 | *B. subtilis* | *trpC2 ansAB::ermC* | LFH → 168 |
| BP234 | *B. subtilis* | *trpC gltP::cat* | Wicke et al., 2019 |
| BP261 | *B. subtilis* | *gltAB::tet* | cDNA GP807 → SP1 |
| BP264 | *B. subtilis* | *amyE::(P<sub>ansAB</sub>-lacZ aphA)* | pBP1110 → SP1 |
| BP265 | *B. subtilis* | *amyE::(P<sub>ansAB</sub>-lacZ aphA) gltAB::tet* | pBP1110 → BP261 |
| BP266 | *B. subtilis* | *ansR::cat* | LFH → SP1 |
| BP267 | *B. subtilis* | *citG::ermC* | LFH → SP1 |
| BP269 | *B. subtilis* | *ansAB::ermC* | cDNA GP1153 → SP1 |
| BP270 | *B. subtilis* | *aspB::spc* | LFH → SP1 |
| BP271 | *B. subtilis* | *amyE::(P<sub>ansAB</sub>-lacZ aphA) ansR::cat* | cDNA BP266 → BP264 |
| BP272 | *B. subtilis* | *amyE::(P<sub>ansAB</sub>-lacZ aphA) citG::ermC* | cDNA BP267 → BP264 |
| BP273 | *B. subtilis* | *amyE::(P<sub>ansAB</sub>-lacZ aphA) ansR::cat gltAB::tet* | cDNA BP266 → BP265 |
| BP274 | *B. subtilis* | *amyE::(P<sub>ansAB</sub>-lacZ aphA) citG::ermC gltAB::tet* | cDNA BP267 → BP265 |
| BP275 | *B. subtilis* SP1 | *amyE::(P<sub>ansAB</sub>-lacZ aphA) ansR::cat citG::ermC* | cDNA BP271 → BP272 |
| BP276 | *B. subtilis* SP1 | *amyE::(P<sub>ansAB</sub>-lacZ aphA) ansR::cat citG::ermC* | cDNA BP273 → BP274 |
| BP279 | *B. subtilis* SP1 | *amyE::(P<sub>ansAB</sub>-lacZ aphA) aspB::spc* | cDNA BP270 → BP264 |
| BP280 | *B. subtilis* SP1 | *gltAB::tet ansAB::ermC* | cDNA BP269 → BP261 |
| BP281 | Suppressor of BP265 | *amyE::(P<sub>ansAB</sub>-lacZ aphA) gltAB::tet ansR +A36 citG +G917* | Selection |
| BP282 | Suppressor of BP265 | *amyE::(P<sub>ansAB</sub>-lacZ aphA) gltAB::tet ansR Δ2456934-2457298* | Selection |
| BP283 | Suppressor of BP265 | *amyE::(P<sub>ansAB</sub>-lacZ aphA) gltAB::tet ansR +A36 citG C1226A* | Selection |
| BP284 | Suppressor of BP265 | *amyE::(P<sub>ansAB</sub>-lacZ aphA) gltAB::tet ansR +A36* | Selection |
| BP285 | Suppressor of BP265 | *amyE::(P<sub>ansAB</sub>-lacZ aphA) gltAB::tet ansR +A36* | Selection |
| BP286 | Suppressor of BP265 | *amyE::(P<sub>ansAB</sub>-lacZ aphA) gltAB::tet ansR +A36* | Selection |
| BP287 | Suppressor of BP265 | *amyE::(P<sub>ansAB</sub>-lacZ aphA) gltAB::tet P<sub>ansAB</sub> (G-10A) citG T876A* | Selection |
| BP288 | Suppressor of BP265 | *amyE::(P<sub>ansAB</sub>-lacZ aphA) gltAB::tet ansR +A36* | Selection |
| BP289 | Suppressor of BP265 | *amyE::(P<sub>ansAB</sub>-lacZ aphA) gltAB::tet ansR +A36 citG Δ3390211-3390276* | Selection |
| BP290 | Suppressor of BP265 | *amyE::(P<sub>ansAB</sub>-lacZ aphA) gltAB::tet ansR +A36* | Selection |
| BP292 | *B. subtilis* | *amyE::(P<sub>ansAB</sub>-lacZ aphA) ansR::cat aspB::spc* | cDNA BP270 → BP271 |
| BP294 | *B. subtilis* | *amyE::(ansR-P<sub>ansR</sub>-lacZ aphA) gltAB::tet* | pBP1111 → BP261 |
| BP296 | Suppressor of BP279 | *amyE::(P<sub>ansAB</sub>-lacZ aphA) aspB::spc citG C686T ansA T43A* | Selection |
| BP297 | Suppressor of BP279 | *amyE::(P<sub>ansAB</sub>-lacZ aphA) aspB::spc citG C312G ansA T43A* | Selection |
| BP298 | Suppressor of BP265 | *amyE::(P<sub>ansAB</sub>-lacZ aphA) gltAB::tet citG A1033T 25.2 kbp amplification including ansAB* | Selection |
| BP364 | Suppressor of BP261 | *gltAB::tet ansR T302C 10.7 kbp deletion including citG* | Selection |
| BP365 | Suppressor of BP261 | *gltAB::tet ansR T169C citG ΔG62* | Selection |
| BP366 | Suppressor of BP279 | *amyE::(P<sub>ansAB</sub>-lacZ aphA) aspB::spc ansR C171A* | Selection |
| BP367 | Suppressor of BP279 | *amyE::(P<sub>ansAB</sub>-lacZ aphA) aspB::spc ansR G3A* | Selection |
| BP368 | Suppressor of BP279 | *amyE::(P<sub>ansAB</sub>-lacZ aphA) aspB::spc ansR G94C* | Selection |
| BP369 | Derivative of BP275 | *amyE::(P<sub>ansAB</sub>-lacZ aphA) ansR::cat citG::ermC gudB ΔG279-C287* | Selection |
| BP370 | Derivatives of BP276 | *amyE::(P<sub>ansAB</sub>-lacZ aphA) gltAB::tet ansR::cat citG::ermC gudB ΔG279-C287* | Selection |
| BP371 | Suppressor of BP275 | *amyE::(P<sub>ansAB</sub>-lacZ aphA) ansR::cat citG::ermC 5.2 kbp amplification including aspB* | Selection |
| BP372 | Suppressor of BP275 | *amyE::(P<sub>ansAB</sub>-lacZ aphA) ansR::cat citG::ermC 5.2 kbp amplification including aspB* | Selection |
| BP373 | Suppressor of BP275 | *amyE::(P<sub>ansAB</sub>-lacZ aphA) ansR::cat citG::ermC rocC C1232T* | Selection |
| BP374 | Suppressor of BP275 | *amyE::(P<sub>ansAB</sub>-lacZ aphA) ansR::cat citG::ermC rocC + T698* | Selection |
| BP375 | Suppressor of BP276 | *amyE::(P<sub>ansAB</sub>-lacZ aphA) gltAB::tet ansR::cat citG::ermC P<sub>dinG</sub> G-63A* | Selection |
| BP376 | Suppressor of BP276 | *amyE::(P<sub>ansAB</sub>-lacZ aphA) gltAB::tet ansR::cat citG::ermC 34.7 kbp amplification including aspB* | Selection |
| BP377 | Suppressor of BP276 | *amyE::(P<sub>ansAB</sub>-lacZ aphA) gltAB::tet ansR::cat citG::ermC odhA ΔT175-A351* | Selection |
| BP378 | Suppressor of BP276 | *amyE::(P<sub>ansAB</sub>-lacZ aphA) gltAB::tet ansR::cat citG::ermC P<sub>odhA</sub> G-189A* | Selection |
| BP382 | *B. subtilis* | *amyE::(ansR-P<sub>ansR</sub>-lacZ aphA)* | pBP1111 → SP1 |
| BP383 | *B. subtilis* | *amyE::(ansR-P<sub>ansR</sub>-lacZ aphA) aspB::spc* | cDNA BP270 → BP382 |
| BP384 | Derivative of BP264 | *amyE::(P<sub>ansAB</sub>-lacZ aphA)* | Selection |
| BP647 | *B. subtilis* | *trpC recN::ermC* | LFH → 168 |
| BP1303 | *B. subtilis* | *trpC gdpP::spc* | Schwedt et al., 2023 |
| XL1-Blue | *E. coli* XL1-Blue | *recA1, endA1, gyrA96, thi-1, hsdR17, supE44, relA1, lac [F' proAB, lacIqZΔM15, Tn10 (Tet<sup>r</sup>)]* | Stratagene |
| Wild type | *C. glutamicum* ATCC13032 | - | Abe et al., 1967 |
| Δ*gdh* Δ*gltB* | *C. glutamicum* ATCC13032 | Δ*gdh* Δ*gltB* | This study |
| Δ*gdh* Δ*gltB* Δ*aspA* | *C. glutamicum* ATCC13032 | Δ*gdh* Δ*gltB* Δ*aspA* | This study |
| Δ*gdh* Δ*gltB* Δ*aspT* | *C. glutamicum* ATCC13032 | Δ*gdh* Δ*gltB* Δ*aspT* | This study |

<sup>a</sup> Arrows indicate strain construction by transformation.

**Table S2.**
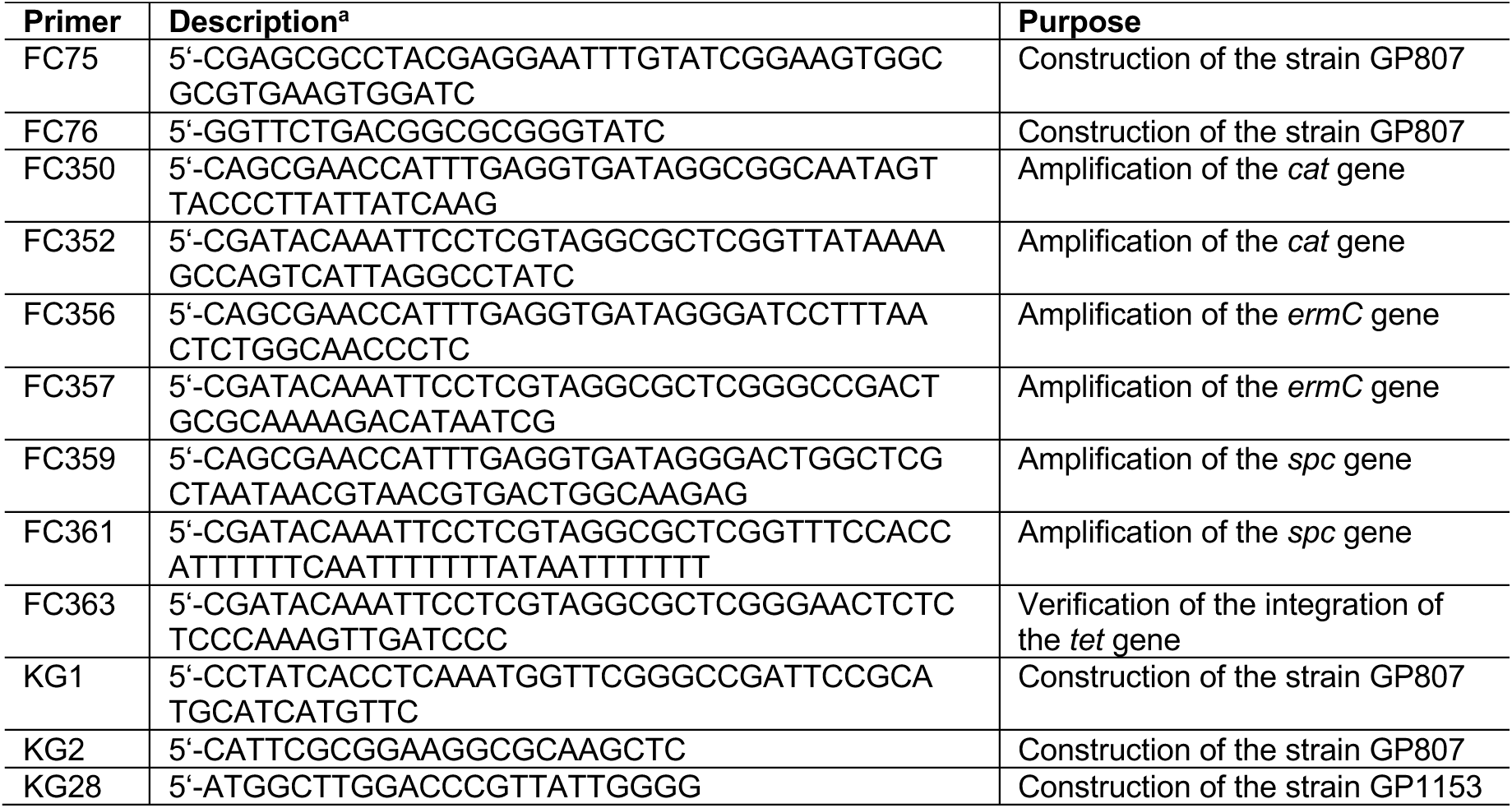
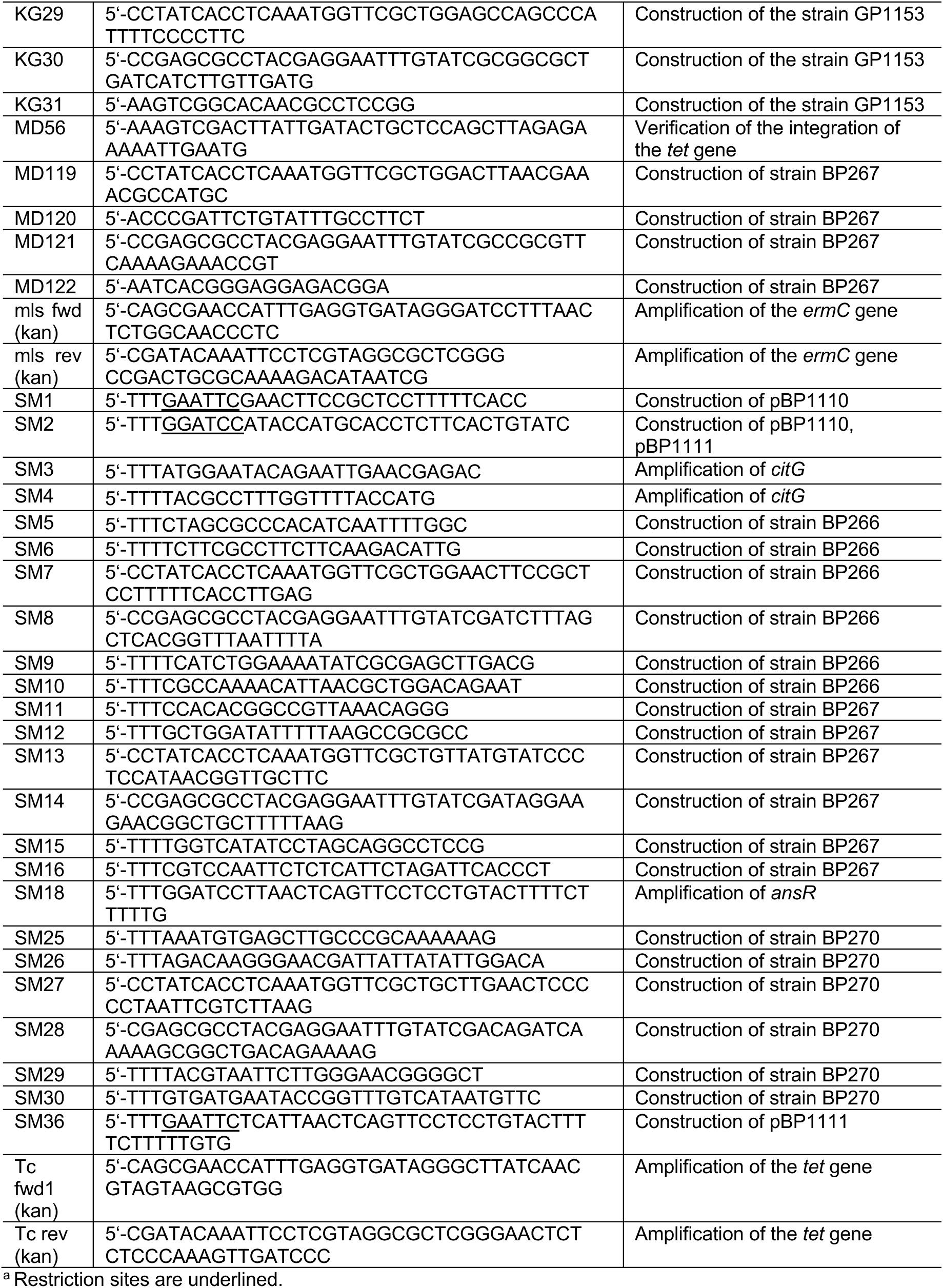
Primers.

**Table S3.** Plasmids.

| Plasmid | Description | Reference, construction |
| --- | --- | --- |
| pAC7 | For the construction of translational *lacZ* fusions, integration into the *amyE* locus | Weinrauch et al., 1991 |
| pBP1110 | pAC7::P<sub>ansAB</sub> | This study |
| pBP1111 | pAC7:: *ansR*-P<sub>ansAB</sub> | This study |
| pDG647 | Template for the amplification of the *ermC* gene | Guérot-Fleury et al., 1995 |
| pDG1514 | Template for the amplification of the *tet* gene | Guérot-Fleury et al., 1995 |
| pDG1726 | Template for the amplification of the *spc* gene | Guérot-Fleury et al., 1995 |
| pGEM-cat | Template for the amplification of the *cat* gene | Laboratory collection |

## Notes

### Competing Interest Statement

The authors have declared no competing interest.

## References

Anagnostopoulos C, Spizizen J 1961. Requirements for transformation in *Bacillus subtilis*. J Bacteriol 81: 741–746.

Barbe V, Cruveiller S, Kunst F, Lenoble P, Meurice G, Sekowska A, Vallenet D, Wang T, Moszer I, Médigue C, Danchin A. 2009. From a consortium sequence to a unified sequence: The *Bacillus subtilis* 168 reference genome a decade later. Microbiology 155: 1758–1775.

Belitsky BR, Sonenshein AL. 1997. Altered transcription activation specificity of a mutant form of *Bacillus subtilis* GltR, a LysR family member. J Bacteriol 179: 1035–1043.

Belitsky BR, Sonenshein AL. 1998. Role and regulation of *Bacillus subtilis* glutamate dehydrogenase genes. J Bacteriol 180: 6298–6305.

Belitsky BR, Sonenshein AL. 2004. Modulation of activity of *Bacillus subtilis* regulatory proteins GltC and TnrA by glutamate dehydrogenase. J Bacteriol 186: 3399–3407.

Belitsky BR, Kim HJ, Sonenshein AL. 2004. CcpA-dependent regulation of *Bacillus subtilis* glutamate dehydrogenase gene expression. J Bacteriol 182: 5939–5947.

Bennett BD, Kimball EH, Gao M, Osterhout R, Van Dien SJ, Rabinowitz JS. 2009. Absolute metabolite concentrations and implied enzyme active site occupancy in *Escherichia coli*. Nat Chem Biol 5: 593–599.

Bohannon DE, Sonenshein AL. 1989. Positive regulation of glutamate biosynthesis in *Bacillus subtilis*. J Bacteriol 171: 4718–4727.

Bremer E, Krämer R 2019. Responses of microorganisms to osmotic stress. Annu Rev Microbiol 73: 313–334.

Brill J, Hoffmann T, Beilsteiner M, Bremer E 2011. Osmotically controlled synthesis of the compatible solute proline is critical for cellular defense of *Bacillus subtilis* against high osmolarity. J Bacteriol 193: 5334–5346.

Calogero S, Gardan R, Glaser P, Schweizer J, Rapoport G, Debarbouillé M 1994. RocR. A novel regulatory protein controlling arginine utilization in *Bacillus subtilis*, belongs to the NtrC/NifA family of transcriptional activators. J Bacteriol 176: 1234–1241.

Choi SK, Saier MH Jr. 2005. Regulation of *sigL* expression by the catabolite control protein CcpA involves roadblock mechanism in *Bacillus subtilis*: potential connection between carbon and nitrogen metabolism. J Bacteriol 187: 6856–6861.

Commichau FM, Forchhammer K, Stülke J. 2006. Regulatory links between carbon and nitrogen metabolism. Curr Opin Microbiol 9: 167–172.

Commichau FM, Wacker I, Schleider J, Blencke HM, Reif I, Tripal P, Stülke J. 2007a. Characterization of *Bacillus subtilis* mutants with carbon source-independent glutamate biosynthesis. J Mol Microbiol Biotechnol 12: 106–113.

Commichau FM, Herzberg C, Tripal P, Valerius O, Stülke J. 2007b. A regulatory protein-protein interaction governs glutamate biosynthesis in *Bacillus subtilis*: the glutamate dehydrogemase RocG moonlights in controlling the transcription factor GltC. Mol Microbiol 65: 642–654.

Commichau FM, Gunka K, Landmann JJ, Stülke J. 2008. Glutamate metabolism in *Bacillus subtilis*: gene expression and enzyme activities evolved to avoid futile cycles and to allow rapid responses to perturbations of the system. J Bacteriol 190: 3557–3567.

Commichau FM, Stülke J. 2008. Trigger enzymes: bifunctional proteins active in metabolism and in controlling gene expression. Mol Microbiol 67: 692–702.

Cotton CA, Bernhardsgrütter I, He H, Burgener S, Schulz L, Paczia N, Dronsella B, Erban A, Toman S, Dempfle M, De Maria A, Kopka J, Lindner SN, Erb TJ, Bar-Even A. 2020. Underground isoleucine biosynthesis pathways in *E. coli*. Elife 9: e54207.

Csonka LN, Ikeda TP, Fletcher SA, Kustu S 1994. The accumulation of glutamate is necessary for optimal growth of *Salmonella typhimurium* in media of high osmolarity but not induction of the *proU* operon. J Bacteriol 176: 6324–6333.

Dormeyer M, Lübke AL, Müller P, Lentes S, Reuß DR, Thürmer A, Stülke j, Daniel R, Brantl, S, Commichau FM. 2017. Hierarchical mutational events compensate for glutamate auxotrophy of a *Bacillus subtilis gltC* mutant. Environ Microbiol Rep 9: 279–289.

Dormeyer M, Lentes S, Richts B, Heermann R, Ischebeck T, Commichau FM. 2019. Variants of the *Bacillus subtilis* LysR-type regulator GltC with altered activator and repressor function. Front Microbiol 10: 2321.

Epstein W 2003. The roles and regulation of potassium in bacteria. Prog Nucleic Acids Res Mol Biol 75: 293–320.

Errington J, Appleby L, Daniel RA, Goodfellow H, Partridge SR, Yudkin MD. 1992. Structure and function of the *spoIIIJ* gene of *Bacillus subtilis*: a vegetatively expressed gene that is essential for sigma G activity at an intermediate stage of sporulation. J Gen Microbiol 138: 2609–2618.

Faires N, Tobisch S, Bachem S, Martin-Verstraete I, Hecker M, Stülke J. 1999. The catabolite control protein CcpA controls ammonium assimilation in *Bacillus subtilis*. J Mol Microbiol Biol 1: 141–148.

Feavers IM, Price V, Moir A. 1988. The regulation of the fumarase (*citG*) gene of *Bacillus subtilis* 168. Mol Gen Genet 211: 465–471.

Fisher SH, Wray LV. 2002. *Bacillus subtilis* 168 contains two differentially regulated genes encoding L-asparaginase. J Bacteriol 184: 2148–2154.

Flórez LA, Gunka K, Polanía R, Tholen S, Stülke J. 2011. SPABBATS: a pathway-discovery method based on Boolean satisfiability that facilitates the characterization of suppressor mutants. BMC Syst Biol 5:5.

Frank C, Hoffmann T, Zelder O, Felle MF, Bremer E. 2021. Enhanced glutamate synthesis and export by the thermotolerant emerging industrial workhorse *Bacillus methanolicus* in response to high osmolarity. Front Microbiol 12: 640980.

Gaballa A, Newton GL, Antelmann H, Parsonage D, Upton H, Rawat M, Claiborne A, Fahey RC, Helmann JD. 2010. Biosynthesis and functions of bacillithiol, a major low-molecular-weight thiol in Bacilli. Proc Nat Acad Sci USA 107: 6482–6486.

Gardan R, Rapoport G, Débarbouillé M. 1995. Expression of the *rocDEF* operon involved in arginine catabolism in *Bacillus subtilis*. J Mol Biol 249:843–856.

Gardan R Rapoport G, Débarbouillé M 1997. Role of the transcriptional activator RocR in the arginine-degradation pathway of *Bacillus subtilis*. Mol Microbiol 24: 825–837.

Guder CJ, Schramm T, Sander T, Link H. 2017. Time-optimized isotope ratio LC-MS/MS for high-throughput quantification of primary metabolites. Anal Chem 89: 1624–1631.

Guérout-Fleury AM, Shazand K, Frandsen N, Stragier P. 1995. Antibiotic resistance cassettes for *Bacillus subtilis*. Gene 167: 335–336.

Gundlach J, Herzberg C, Kaever V, Gunka K, Hoffmann T, Weiß M, Gibhardt J, Thürmer A, Hertel D, Daniel R, Bremer E, Commichau FM, Stülke J. 2017. Control of potassium homeostasis is an essential function of the second messenger cyclic di-AMP in *Bacillus subtilis*. Sci Signal 10: eaal3011.

Gundlach J, Commichau FM, Stülke J. 2018. Perspective of ions and messengers: an intricate link between potassium, glutamate, and cyclic di-AMP. Curr Genet 64: 191–195.

Gunka K, Newman JA, Commichau FM, Herzberg C, Rodrigues C, Hewitt L, Lewis RJ, Stülke J. 2010. Functional dissection of a trigger enzyme: mutations of the *Bacillus subtilis* glutamate dehydrogenase RocG that affect differentially its catalytic activity and regulatory properties. J Mol Biol 400: 815–827.

Gunka K, Tholen S, Gerwig J, Herzberg C, Stülke J, Commichau FM. 2012. A high-frequency mutation in *Bacillus subtilis*: requirements for the decryptification of the *gudB* glutamate dehydrogenase gene. J Bacteriol 194: 1036–1044.

Gunka K, Commichau FM. 2012. Control of glutamate homeostasis in *Bacillus subtilis*: a complex interplay between ammonium assimilation, glutamate biosynthesis and degradation. Mol Microbiol 85: 213–224.

Gunka K, Stannek L, Care RA, Commichau FM. 2013. Selection-driven accumulation of suppressor mutants in *Bacillus subtilis*: the apparent high mutation frequency of the cryptic *gudB* gene and the rapid clonal expansion of *gudB*(+) suppressors are due to growth under selection. PLoS One 8: e66120.

Hartmann MD. 2022. A complex struggle for direction. Nat Chem Biol 18: 119–120.

Helling RB. 1994. Why does *Escherichia coli* have two primary pathways for synthesis of glutamate? J Bacteriol 176: 4664–4668.

Helling RB. 1998. Pathway choice in glutamate synthesis in *Escherichia coli*. J Bacteriol 180: 4571–4575.

HoffmannT, Bleisteiner M, Sappa PK, Steil L, Mäder U, Völker U, Bremer E 2017. Synthesis of the compatible solute proline by *Bacillus subtilis*: point mutations rendering the osmotically controlled *proHJ* promoter hyperactive. Environ Microbiol 19: 3700–3720.

Hudson RC, Daniel RM 1993. L-Glutamate dehydrogenases: distribution, properties and mechanism. Comp Biochem Physiol B. 106: 767–792.

Huerta-Cepas J, Serra F, Bork P (2016) ETE 3: Reconstruction, analysis, and visualization of phylogenomic data. Mol Biol Evol 33: 1635–1638.

Iacometti C, Marx K, Hönick M, Biletskaia V, Schulz-Mirbach H, Dronsella B, Satanowski A, Delmas VA, Berger A, Dubois I, Bouzon M, Döring V, Noor E, Bar-Even A, Lindner S 2022. Activating silent glycolytic bypasses in *Escherichia coli*. Biodes Res. 2022:9859643.

Ikeda TP, Shauger AE, Kustu S. 1996. *Salmonella typhimurium* apparently perceives external nitrogen limitation as internal glutamine limitation. J Mol Biol 259: 589–607.

Jayaraman V, Lee DJ, Elad N, Vimer S, Sharon M, Fraser JS, Tawfik DS. 2022. A counter-enzyme complex regulates glutamate metabolism in *Bacillus subtilis*. Nat Chem Biol 18: 161–170.

Kearse M, Moah R, Wilson A, Stones-Havas S, Cheung M, Sturrock S, Buxton S, Cooper A, Markowitz S, Duran C, Thierer T, Asthon B, Meintjes P, Drummond A (2012) Geneious Basic: An integrated and extendable desktop software platform for the organization and analysis of sequence data. Bioinformatics 28: 1647–1649.

Keilhauer C, Eggeling L, Sahm H. 1993. Isoleucine synthesis in *Corynebacterium glutamicum*: moleculare analysis of the *ilvB-ilvN-ilvC* operon. J Bacteriol 175: 5595–5603.

Kim J, Copley SD. 2007. Why metabolic enzymes are essential or non-essential for growth of *Escherichia coli* K12 on glucose. Biochemistry 46: 12501–12511.

Kim J, Kershner JP, Novikov Y, Shoemaker RK, Copley SD. 2010. Three serendipitous pathways in *E. coli* can bypass a block in pyridoxal-5’-phosphate synthesis. Mol Syst Biol 6: 436.

Kim J, Flood JJ, Kristofich MR, Gidfar C, Morgenthaler AB, Fuhrer T, Sauer U, Snyder D, Cooper VS, Ebmeier CC, Old WM, Copley SD. 2019. Hidden resources in the *E. coli* genome restore PLP synthesis and robust growth after deletion of the essential gene *pdxB*. Proc Natl Acad Sci USA 116: 24164–24173.

Krüger L, Herzberg C, Warneke R, Poehlein A, Stautz J, Weiß M, Daniel R, Hänelt I, Stülke J. 2020. Two ways to convert a low-affinity potassium channel to high affinity: control of *Bacillus subtilis* KtrCD by glutamate. J Bacteriol 202: e00138–20.

Krüger L, Herzberg C, Rath H, Pedreira T, Ischebeck T, Poehlein A, Gundlach J, Daniel R, Völker U, Mäder U, Stülke J. 2021. Essentiality of c-di-AMP in *Bacillus subtilis*: bypassing mutations converge in potassium and glutamate homeostasis. PLoS Genet 17: e1009092.

Kumada Y, Benson DR, Hillemann D, Hosted TJ, Rochefort DA, Thompson CJ, Wohlleben W, Tateno Y. 1993. Evolution of the glutamine synthetase gene, one of the oldest existing and functioning genes. Proc Natl Acad Sci USA 90: 3009–3013.

Kunst F, Rapoport G. 1995. Salt stress is an environmental signal affecting degradative enzyme synthesis in *Bacillus subtilis*. J Bacteriol 177: 2403–2407.

Larsson JT, Rogstam A, von Wachenfeld C 200. Coordinated patterns of cytochrome bd and lactate dehydrogenase expression in *Bacillus subtilis*. Microbiology 151: 3323–3335.

Liu H, Jeffery CJ. 2020. Moonlighting proteins in the fuzzy logic of cellular metabolism. Molecules 25: 3440.

Magasanik B. 1996. Regulation of nitrogen utilization, p. 1344-1356. In FC Neidhardt et al. (ed), Escherichia coli and Salmonella typhimurium: cellular and molecular biology, vol. 1. ASM Press, Washington DC.

Magasanik B. 2003. Ammonia assimilation by *Saccharomyces cerevisiae*. Eukaryot Cell 2: 827–829.

McLaggan D, Naprstek J, Buurman ET, Epstein W 1994. Interdependence of K+ and glutamate accumulation during osmotic adaptation of *Escherichia coli*. J Biol Chem 269: 1911–1917.

Miller CM, Baumberg S, Stockley PG. 1997. Operator interactions by the *Bacillus subtilis* arginine repressor/activator, AhrC: a novel positioning and DNA-mediated assembly of a transcriptional activator at catabolic sites. Mol Microbiol 26: 37–46.

Moir A, Feavers IM, Guest JR. 1984. Characterization of the fumarase gene of *Bacillus subtilis* 168 cloned and expressed in *Escherichia coli* K12. J Gen Microbiol 130: 3009–3017.

Miles JS, Guest JR. 1985. Complete nucleotide sequence of the fumarase gene (*citG*) of *Bacillus subtilis* 168. Nucleic Acids Res 13: 131–140.

Noda-Garcia L, Romero Romero ML, Longo LM, Kolodkin-Gal I, Tawfik DS. 2017. *Bacilli* glutamate dehydrogenases diverged via coevolution of transcription and enzyme regulation. EMBO Rep 18: 1139–1149.

Oh YK, Palsson B, Park SM, Schilling CH, Mahadevan R. 2007. Genome-scale reconstruction of metabolic network in *Bacillus subtilis* based on high-throughput phenotyping and gene essentiality data. J Biol Chem 282: 28791–28799.

Park JO, Rubin SA, Xu YF, Amador-Noguez D, Fan J, Shlomi T, Rabinowitz JD. 2016. Metabolite concentrations, fluxes and free energies imply efficient enzyme usage. Nat Chem Biol 12: 482–489.

Parsot C. 1986. Evolution of biosynthetic pathways: a common ancestor for threonine synthase, threonine dehydratase and D-serine dehydratase. Embo J 5: 3013–3019.

Picossi S, Belitsky BR, Sonenshein AL. 2007. Molecular mechanism of the regulation of *Bacillus subtilis gltAB* expression by GltC. J Mol Biol 365:1298–1313.

Pontrelli S, Fricke RCB, Teoh ST, Lavina WA, Putri AP, Fitz-Gibbon S, Chung M, Pellegrini M, Fukusaki E, Liao JC. 2018. Metabolic repair through emergence of new pathways in *Escherichia coli*. Nat Chem Biol 14: 1005–1009.

Quinn CL, Stephenson BT, Switzer RL 1991. Functional organization and nucleotide sequence of the *Bacillus subtilis* pyrimidine biosynthetic operon. J Bacteriol 266: 9113–9127.

Rehm N, Georgi T, Hiery E, Degner Schmiedl A, Burkovski A, Bott M. 2010. L-Glutamine as nitrogen source for *Corynebacterium glutamicum*: derepression of the AmtR regulon and implications for nitrogen sensing. Microbiology (Reading) 156: 3180–3193.

Reizer LJ. 2003. Nitrogen assimilation and global regulation in *Escherichia coli*. Ann Rev Microbiol 57: 155–176.

Reuß DR, Rath H, Thürmer A, Benda M, Daniel R, Völker U, Mäder U, Commichau FM, Stülke J. 2018. Changes of DNA topology affect the global transcription landscape and allow rapid growth of a *Bacillus subtilis* mutant lacking carbon catabolite repression. Metab Eng 45: 171–179.

Richts B, Commichau FM. Underground metabolism facilitates the evolution of novel pathways for vitamin B6 biosynthesis. Appl Microbiol Biotechnol 105: 2297–2305.

Richts B, Lentes S, Poehlein A, Daniel R, Commichau FM. 2021. A *Bacillus subtilis pdxT* mutant suppresses vitamin B6 limitation by acquiring mutations enhancing *pdxS* gene dosage and ammonium assimilation. Environ Microbiol Rep 13: 218–233.

Rosenberg J, Müller P, Lentes S, Thiele MJ, Zeigler DR, Tödter D, Paulus, H, Brantl S, Stülke J, Commichau FM. 2016. ThrR, a DNA-binding transcription factor involved in controlling threonine biosynthesis in *Bacillus subtilis*. Mol Microbiol. 101: 879–893.

Rosenberg J, Yeak KC, Commichau FM. 2018 A two-step evolutionary process establishes a non-native vitamin B6 pathway in *Bacillus subtilis*. Environ Microbiol 20: 156–168.

Rosenberg J, Commichau FM. 2019. Harnessing underground metabolism for pathway development. Trends Biotechnol 37: 29–37.

Rosenow EC. 1919. Studies on elective localization focal infection with special reference to oral sepsis. J Dental Res 1: 205–267.

Saum SH, Sydow JF, Palm P, Pfeiffer F, Oesterheld D, Müller V 2006. Biochemical and molecular characterization of the biosynthesis of glutamine and glutamate, two major compatible solutes in the moderately halophilic bacterium *Halobacillus halophilus*. J Bacteriol 188: 6808–6815.

Schujman GE, Paoletti L, Grossmann AD, de Mendoza D 2003. FapR, a bacterial transcription factor involved in global regulation of membrane lipid biosynthesis. Dev Cell 4: 663–672.

Schulz-Mirbach H, Müller A, Wu T, Pfister P, Aslan S, Schada von Borzyskowski A, Erb TJ, Bar-Even A, Lindner SN. 2022. On the flexibility of the cellular amination network in *E coli*. Elife 11: e77492.

Schwedt I, Schöne K, Eckert M, Pizzinato M, Winkler L, Knotkova B, Richts B, Hau JL, Steuber J, Mireles R, Noda-Garcia L, Fritz G, Mittelstädt C, Hertel R, Commichau FM 2023. The low mutational flexibility of the 5-enolpyruvyl-shikimate-3-phosphate synthase in *Bacillus subtilis* is due to a higher demand for shikimate pathway intermediates. Environ Microbiol In press.

Sezonov G, Joseleau-Petit D, D’Ari R 2007. *Escherichia coli* physiology in Luria-Bertani broth. J Bacteriol 8746–8749.

Sonenshein AL. 2007. Control of key metabolic intersections in *Bacillus subtilis*. Nat Rev Microbiol 5: 917–927.

Stannek L, Thiele MJ, Ischebeck T, Gunka K, Hammer E, Völker U, Commichau FM. 2015. Evidence for synergistic control of glutamate biosynthesis by glutamate dehydrogenases and glutamate in *Bacillus subtilis*. Environ Microbiol 17: 3379–3390.

Stecker D, Hoffmann T, Link H, Commichau FM, Bremer E 2022. L-Proline synthesis mutants of *Bacillus subtilis* overcome osmotic sensitivity by genetically adapting L-arginine metabolism. Front Microbiol 13: 908304.

Sun DX, Setlow P. 1991. Cloning, nucleotide sequence, and expression of the *Bacillus subtilis ans* operon, which codes for L-asparaginase and L-asparase. J Bacteriol 173: 3831–3845.

Sun D, Setlow P. 1993. Cloning and nucleotide sequence of the *Bacillus subtilis ansR* gene, which encodes a repressor of the *ans* operon coding for L-asparaginase and L-aspartase. J Bacteriol 175: 2501–2506.

Suzuki A, Knaff DB. 2005. Glutamate synthase: structural, mechanistic and regulatory properties, and role in the amino acid metabolism. Photosynth Res 83: 191–217.

Veeravalli K, Boyd D, Iverson BL, Beckwith J Georgiou G. 2011. Laboratory evolution of glutathione biosynthesis reveals natural compensatory pathways. Nat Chem Biol 7: 101–105.

Viola RE. 2000. L-aspartase: new tricks from an old enzyme. Adv Enzymol Relat Areas Mol Biol. 74: 295–341.

Wacker I, Ludwig H, Reif I, Blencke HM, Detsch C, Stülke J. 2003. The regulatory link between carbon and nitrogen metabolism in *Bacillus subtilis*: regulation of the *gltAB* operon by the carbon catabolite control protein CcpA. Microbiology 149: 3001–3009.

Warneke R, Garbers TB, Herzberg C, Aschenbrandt G, Ficner R, Stülke J 2023. Ornithine is the central intermediate in the arginine degradative pathway and its regulation in *Bacillus subtilis*. J Biol Chem 299: 104944.

Weinrauch Y, Penchev R, Dubnau E, Smith I, Dubnau D 1990. A *Bacillus subtilis* regulatory gene product for genetic competence and sporulation resembles sensor protein members of the bacterial two-component signal-transduction systems. Genes Dev 4: 860–872.

Wicke D, Schulz LM, Lentes S, Scholz P, Poehlein A, Gibhardt J, Daniel R, Ischebeck T, Commichau FM. 2019. Identification of the first glyphosate transporter by genomic adaptation. Environ Microbiol 21: 1287–1305.

Widderich N, Rodrigues CD, Commichau FM, Fischer KE, Ramirez-Guadiana FH, Rudner DZ, Bremer E. 2016. Salt-sensitivity of σ(H) and Spo0A prevents sporulation of *Bacillus subtilis* at high osmolarity avoiding death during cellular differentiation. Mol Microbiol 100: 108–124.

Wohlheuter RM, Schutt H, Holzer H. 1973. Regulation of glutamine synthesis in vivo in E. coli. In The Enzymes of Glutamine Metabolism. Prusiner SB, Stadtmann ER (eds). New York: Academic Press, pp. 45–64.

Zaprasis A, Brill J, Thüring M, Wünsche G, Heun M, Barzantny H, Hoffmann T, Bremer E 2013. Osmoprotection of *Bacillus subtilis* through import and proteolysis of proline-containing peptides. Appl Environ Microbiol 79: 576–587.

Zaprasis A, Hoffmann T, Wünsche G, Flórez LA, Stülke J, Bremer E. 2014. Mutational activation of the RocR activator and of a cryptic *rocDEF* promoter bypass loss of the initial steps of proline biosynthesis in *Bacillus subtilis*. Environ Microbiol 16: 701–717.

Zaprasis A, Bleisteiner M, Kerres A, Hoffmann T, Bremer E. 2015. Uptake of amino acids and their metabolic conversion into the compatible solute proline confers osmoprotection to *Bacillus subtilis*. Appl Environ Microbiol 81: 250–259.

Zeigler DR, Prágai Z, Rodriguez S, Chevreux B. Muffler A, Albert T, Bai R, Wyss M, Perkins JB. 2008. The origins of 168, W23, and other *Bacillus subtilis* legacy strains. J Bacteriol 190: 6983–6995.

Zhao H, Roistacher DM, Helmann JD. 2018. Aspartate deficiency limits peptidoglycan synthesis and sensitizes cells to antibiotics targeting cell wall synthesis in *Bacillus subtilis*. Mol Microbiol 109: 826–844.

## Supporting information references

Abe S, Takayama K, Kinoshita S (1967) Taxonomical studies on glutamic acid-producing bacteria. J Gen Appl Microbiol 13: 279–301.

Blasco B, Stenta M, Alonso-Sarduy L, Dietler G, Peraro MD, Cole ST, Pojer F. 2011. Atypical DNA recognition mechanism used by the EspR virulence regulator of *Mycobacterium tuberculosis*. Mol Microbiol 82: 251–264.

Guo J, Coker AR, Wood SP, Cooper JB, Chohan SM, Rashi N, Akhtar M. 2017. Structure and function of the thermostable L-asparaginase from *Thermococcus kodakarensis*. Acta Crystallogr D Struct Biol 73: 889–895.

Meyer P, Evrin C, Briozzo P, Joly N, Barzu O, Gilles AM. 2008. Structural and functional characterization of *Escherichia coli* UMP kinase in complex with its allosteric regulator GTP. J Biol Chem 283: 36011.

Omasits U, Ahrens CH, Müller S, Wollscheid B. 2014. Protter: interactive protein feature visualization and integration with experimental proteomic data. Bioinformatics 30: 884–886.

Parris KD, Lin L, Tam A, Mathew R, Hixon J, Stahl M, Fritz CC, Seehra J, Somers WS. 2000. Crystal structures of substrate binding to *Bacillus subtilis* holo-(acyl carrier protein) synthase reveal a novel trimeric arrangement of molecules resulting in three active sites. Structure 8: 883–895.

Rehm N, Georgi T, Hiery E, Degner U, Schmiedl A, Burkovski A, Bott M (2010) L-Glutamine as a nitrogen source for *Corynebacterium glutamicum*: derepression of the AmtR regulon and implications for nitrogen sensing. Microbiology 156: 3180–3193.

Richts B, Hertel R, Potot S, Poehlein A, Daniel R, Schyns G, Prágai Z, Commichau FM (2020) Complete genome sequence of the prototrophic *Bacillus subtilis* subsp. *subtilis* strain SP1. Microbiol Resourc Announc 9: e00825–20.

Schwedt I, Schöne K, Eckert M, Pizzinato M, Winkler L, Knotkova B, Richts B, Hau JL, Steuber J, Mireles R, Noda-Garcia L, Fritz G, Mittelstädt C, Hertel R, Commichau FM (2023) The low mutational flexibility of the EPSP synthase in *Bacillus subtilis* is due to a higher demand for shikimate pathway intermediates. Environ Microbiol. doi: 10.1111/1462-2920.16518.

Waterhouse A, Bertoni M, Bienert S, Studer G, Tauriello G, Gumienny R et al. 2018. SWISS-MODEL: homology modeling of protein structures and complexes. Nucleic Acids Res 46: W296–W303.

Weinrauch Y, Msadek T, Kunst F, Dubnau D (1991) Sequence and properties of *comQ*, a new competence regulatory gene of *Bacillus subtilis*. J Bacteriol 173: 5685–5693.

Wicke D, Schulz LM, Lentes S, Scholz P, Poehlein A, Gibhardt J, Daniel R, Ischebeck T, Commichau FM (2019) Identification of the first glyphosate transporter by genomic adaptation. Environ Microbiol 21: 1287–1305.

